# The diverse evolutionary histories of domesticated metaviral capsid genes in mammals

**DOI:** 10.1101/2023.09.17.558119

**Authors:** William S. Henriques, Janet M. Young, Artem Nemudryi, Anna Nemudraia, Blake Wiedenheft, Harmit S. Malik

## Abstract

Selfish genetic elements and their remnants comprise at least half of the human genome. Active transposons duplicate by inserting copies at new sites in a host genome. Following insertion, transposons can acquire mutations that render them inactive; the accrual of additional mutations can render them unrecognizable over time. However, in rare instances, segments of transposons become useful for the host, in a process called gene domestication. Using the first complete human genome assembly and 25 additional vertebrate genomes, we analyzed the evolutionary trajectories and functional potential of genes domesticated from the capsid genes of *Metaviridae*, a retroviral-like retrotransposon family. Our analysis reveals four families of domesticated capsid genes in placental mammals with varied evolutionary outcomes, ranging from universal retention to lineage-specific duplications or losses and from purifying selection to lineage-specific rapid evolution. The four families of domesticated capsid genes have divergent amino-terminal domains, inherited from four distinct ancestral metaviruses. Structural predictions reveal that many domesticated genes encode a previously unrecognized RNA-binding domain retained in multiple paralogs in mammalian genomes both adjacent to and independent from the capsid domain. Collectively, our study reveals diverse outcomes of domestication of diverse metaviruses, which led to structurally and evolutionarily diverse genes that encode important, but still largely-unknown functions in placental mammals.

## INTRODUCTION

At least half of the human genome is derived from selfish genetic elements (Smit 1999; Venter et al. 2001; Li et al. 2001; de Koning et al. 2011). Transposons are selfish genetic elements that encode key proteins needed to replicate and spread independent of host control (Dawkins 1976; Doolittle and Sapienza 1980; Orgel and Crick 1980). In the human genome, a small number of transposons retain the capacity to ‘jump’ to new regions of the genome, whereas the majority are found immobilized and in various stages of degradation (Venter et al. 2001; Deininger and Batzer 2002). New transposon insertions can be harmful; they can interrupt host genes, alter the expression patterns of adjacent genes, or lead to ectopic recombination (Boissinot et al. 2006). Although some new insertion events in somatic cells are implicated in disease (Burns 2017; Burns 2020), most insertions are functionally inconsequential (Baillie et al. 2011; Fueyo et al. 2022).

Although somatic insertions in multicellular organisms are an evolutionary ‘dead-end’, transposon insertions in the germline are heritable. These insertions become a substrate for natural selection. Insertions that are deleterious to host fitness are rapidly eliminated from the population by purifying selection, whereas insertions that are inconsequential accumulate mutations at rates consistent with genomic mutation rates (reviewed in (Mager and Stoye 2015; Wells and Feschotte 2020). However, on rare occasions, pieces of mutationally decaying transposons acquire host functions and are subsequently protected from mutational abrasion by a process known as gene domestication, exaptation, or co-option (J. Brandt et al. 2005; Modzelewski et al. 2022). After acquiring host function, such transposon-derived regions are no longer capable of autonomous replication and evolve just as any other host gene subject to purifying selection.

In mammalian genomes, the most common class of selfish elements are retrotransposons, which generate new copies using a ‘copy-and-paste’ mechanism (Burns and Boeke 2012; Finnegan 2012; Krupovic et al. 2018; Dodonova et al. 2019). Retrotransposons in the human genome can be classified into two broad groups: Non-long-terminal-repeat (non-LTR) Line1 retrotransposons (and non-autonomous Alu and SVA elements that rely on Line1 machinery) or LTR retrotransposons, which include endogenous retroviruses (ERVs)(Cordaux and Batzer 2009), as well as four other families of LTR retrotransposon from the order of reverse-transcribing viruses, *Ortervirales: Metaviridae* (formerly Ty3/Gypsy)*, Caulimoviridate, Belpaoviridae,* and *Pseudoviridae* (formerly Ty1) *(Krupovic et al. 2018)*. LTR retrotransposons encode structural group-specific antigen (*gag)* genes and enzymatic polymerase (*pol*) genes, flanked by two LTRs. Their ‘exogenous’ (transmissible) counterparts also encode an envelope protein (*env*) (Doolittle et al. 1989; Hayward 2017), which mediates membrane fusion needed for infection, and can subvert immune defenses via an immunosuppressive domain (Ashkenazi et al. 2013).

ERV genes have been repeatedly domesticated in mammals for diverse host functions, including reproduction and viral defense. The *syncytin* env-like genes of placental mammals present a spectacular example of domestication, mediating membrane fusion events needed to form multinucleated syncytial trophoblast cells in the placenta (Kim et al. 2004; Dupressoir et al. 2012). *Syncytin* gene domestication has occurred recurrently in a remarkable example of convergence, with at least seven independent events in different mammalian orders (Dupressoir et al. 2012; Lavialle et al. 2013), and in unusual lineages of lizard (Cornelis et al. 2017) and fish (Henzy et al. 2017). In addition to domestication for placental function, multiple retroviral envelope genes have also been domesticated for retroviral defense in mice (Ikeda et al. 1985; Taylor et al. 2001), humans (Frank et al. 2022), sheep (Varela et al. 2009), cats (Ito et al. 2013; Ito et al. 2016), and chickens (Robinson et al. 1981). These co-opted envelope genes reveal two key ideas about gene domestication. First, the same viral protein can be independently repurposed for similar functions in different hosts. Second, domesticated copies of a single viral domain can serve diverse functions even within a single host.

The *gag* genes of reverse-transcribing viruses have been best studied in retroviruses such as HIV-1, and to a lesser extent in *Metaviridae* such as Ty3. Gag genes encode a polyprotein that includes capsid, matrix, and nucleocapsid domains that together package the viral genome during the virion assembly process (Dodonova et al. 2019; Olson and Musier-Forsyth 2019). Like *env*, *gag* has been domesticated for retroviral restriction in mice as the *Fv1* gene (Bénit et al. 1997: 1; Yap et al. 2014: 1; Young et al. 2018: 1), but also for other critical host functions (J. Brandt et al. 2005; Campillos et al. 2006; Ono et al. 2006; Sekita et al. 2008; Kokošar and Kordiš 2013). At least four domesticated capsid genes have been shown to assemble into capsid-like structures (Pastuzyn et al. 2018; Abed et al. 2019; Erlendsson et al. 2019; Segel et al. 2021; Xu et al. 2023), which perform essential functions in host reproduction (Ono et al. 2006) and neuronal function (Nikolaienko et al. 2018; Pastuzyn et al. 2018). One of these *gag-*derived genes is vertebrate *Arc (A*ctivity-*R*egulated, *C*ytoskeletal-associated), which originated from an ancient retroelement from the clade *Metaviridae* (formerly known as Ty3/Gypsy). ARC protein forms capsid-like structures and functions in the brain to regulate learning and memory as a signaling hub and messenger RNA shuttle in neurons (Nikolaienko et al. 2018; Pastuzyn et al. 2018). An independent domestication event from a distinct *Metaviridae* lineage led to *dArc1* in *Drosophila* species, which also forms capsid-like structures and packages mRNA for intercellular neuronal signaling (Ashley et al. 2018; Erlendsson et al. 2020). Together, *Arc* and *dArc1* elegantly demonstrate the functional convergence of independently domesticated genes in animal lineages. Although other domesticated *gag* genes also encode proteins capable of assembling into capsid-like structures, most remain functionally uncharacterized (Xu et al. 2023).

Both independent ancient *Metaviridae* domestication events and post-domestication duplications gave rise to over two dozen domesticated *gag*-like genes in the human genome, including *ARC* (Campillos et al. 2006; Kokošar and Kordiš 2013) and two small gene families, often referred to as the *PNMA* (Paraneoplastic Ma antigens) and *SIRH/RTL* (Sushi-Ichi-related Retrotransposon-Homolog / RetroTransposon-Like) families (J. Brandt et al. 2005; Campillos et al. 2006; Kaneko-Ishino and Ishino 2015). Among the best characterized of these genes are *Peg10* (Paternally Expressed Gene 10, a SIRH/RTL family member) and *Rtl1* (retrotransposon-like 1) (Butler et al. 2001; Charlier et al. 2001; Ono et al. 2001), both of which encode proteins that contain capsid domains and are essential for successful embryonic development (Ono et al. 2006; Sekita et al. 2008; Segel et al. 2021). Like ARC, PEG10 viral-like particles (VLPs) can package nucleic acid; PEG10 VLPs have been repurposed as delivery vehicles for custom nucleic acid cargos with potential therapeutic applications (Clark et al. 2007; Abed et al. 2019; Segel et al. 2021). While roles are emerging for some *PNMA/SIRH* genes (Foley et al. 2008; Irie et al. 2015; Lee et al. 2016; Abed et al. 2019; Irie et al. 2021; Ishino et al. 2023), most remain functionally uncharacterized. Similarly, in-depth evolutionary characterization has only been performed for a few of these genes, revealing a mixture of retention and lineage-specific gene loss (J. Brandt et al. 2005; Irie et al. 2015; Irie et al. 2021). Gene loss in some species is surprising given the essentiality of orthologs of these genes in mice.

Given the important biological functions of domesticated capsid-like human genes, and their therapeutic potential, we sought a deeper structural and evolutionary understanding of these genes, building on previous evolutionary surveys across the family (Campillos et al. 2006; Kokošar and Kordiš 2013), or deeper surveys of individual genes (J. Brandt et al. 2005; Irie et al. 2015; Irie et al. 2021). Seeking a deeper evolutionary synthesis, we took advantage of state-of-the-art tools for structure prediction and homology detection - AlphaFold, Foldseek, and DALI (Holm 2020; Jumper et al. 2021; Kempen et al. 2022) - and the availability of numerous mammalian genome assemblies to carry out a systematic bioinformatic study of the architecture, evolution, expression, and predicted protein structure of capsid-derived sequences. Focusing on the first telomere-to-telomere assembly of the human genome and 25 additional vertebrate genomes, we found that most of the approximately two dozen *bona fide* domesticated metaviral capsid-like genes show clear signatures of purifying selection. About half of these have been strictly retained across placental mammals, suggesting functions common to all lineages. Other genes have undergone lineage-specific gene duplication, pseudogenization, or loss, implying lineage-specific retention or loss of functions. For a small subset of domesticated genes, we find evidence of positive selection, indicating their involvement in as-yet-undiscovered but ongoing genetic conflicts or acquisition of novel functions. Our re-examination of the domain architecture finds no evidence for a canonical retroviral matrix-like domain neighboring the capsid in either the domesticated capsid genes or their active metaviral relatives. Instead, we find that amino-terminal domains (NTD) are widely divergent between domesticated capsid gene families, reflecting their domestication from four distinct *Metaviridae* families with distinct domain architectures. These divergent NTDs include a previously unnoticed RNA-binding domain (RBD) encoded by the *PNMA* family, which has undergone dramatic expansion in mammalian genomes and can be retained either with or without an associated capsid domain. Our study reveals that recurrent domestication of the *gag* domain from structurally diverse *Metaviridae* gave rise to genes with distinct evolutionary trajectories in placental mammals.

## RESULTS

### Divergent metaviral capsid-derived genes in the human genome

Previous analyses identified 85 human genes with homology to retroviral- or retrotransposon-encoded *gag* genes (Campillos et al. 2006), demonstrating that *gag* genes have been recurrently repurposed for host function. We wanted to update this analysis by searching the newly available and complete Telomere-to-Telomere (T2T) human genome assembly (including the Y chromosome) (Nurk et al. 2021; Rhie et al. 2022). We focused on the capsid domain of the *gag* gene, which shares homology across reverse transcribing viruses (Krupovic and Koonin 2017a). To ensure capture of remote homologs, we built Hidden Markov Models (HMMs) guided by atomic structures and AlphaFold predictions of retroviral and retrotransposon capsid domains. We generated a separate HMM for each of the three clades of endogenous retroviruses (*Orthoretrovirinae)* as well as *Spumavirinae* (Gifford et al. 2018) and the three major clades of LTR retrotransposons found in vertebrates: *Metaviridae* (previously known as Ty3/gypsy), *Pseudoviridae* (previously Ty1/copia), and *Belpaoviridae* (Krupovic and Koonin 2017a; Krupovic et al. 2018) *(See Methods,* **Figure S1, Table S1**).

We found 24 of the 85 metaviral*-*derived capsid genes than previously identified (Campillos et al. 2006). The major reason for this disparity is that the previous study contained SCAN domains, which have been proposed to originate from ancient retrotransposon *gag* domains (Ivanov et al. 2005; Emerson and Thomas 2011; Kaneko-Ishino 2012; Krupovic and Koonin 2017b). However, none of our HMMs for the capsid domain detected the numerous known SCAN-domain-containing proteins in the human genome. To keep our analysis focused, we did not build separate HMMs for the SCAN domain. Querying a six-open reading frame translation (between stop codons) of the T2T assembly with our custom capsid-specific HMMs, we identified 3,140 discrete capsid-like sequences from *Orthoretrovirinae* and *Metaviridae*, but none from *Pseudoviridae*, *Belpaoviridae,* or *Spumavirinae,* consistent with previous reports (Sperber et al. 2007; Blikstad et al. 2008).

Of the 3,140 discrete capsid-like sequences, we found only 467 capsid sequences that contain both a start codon and have the potential to encode a full-length capsid domain, whereas the remaining 2,673 contain premature stop codons or frameshift mutations and are unlikely to encode functional proteins. To understand the relationship between these 467 sequences, we aligned them to each other. This alignment allowed us to identify and remove poorly aligning sequences likely due to internal deletions, ultimately yielding 212 well-aligned open-reading frames that encode a full-length capsid domain. We used this alignment to construct an unrooted maximum-likelihood phylogenetic tree (**Figure 1**).

**Figure 1.**
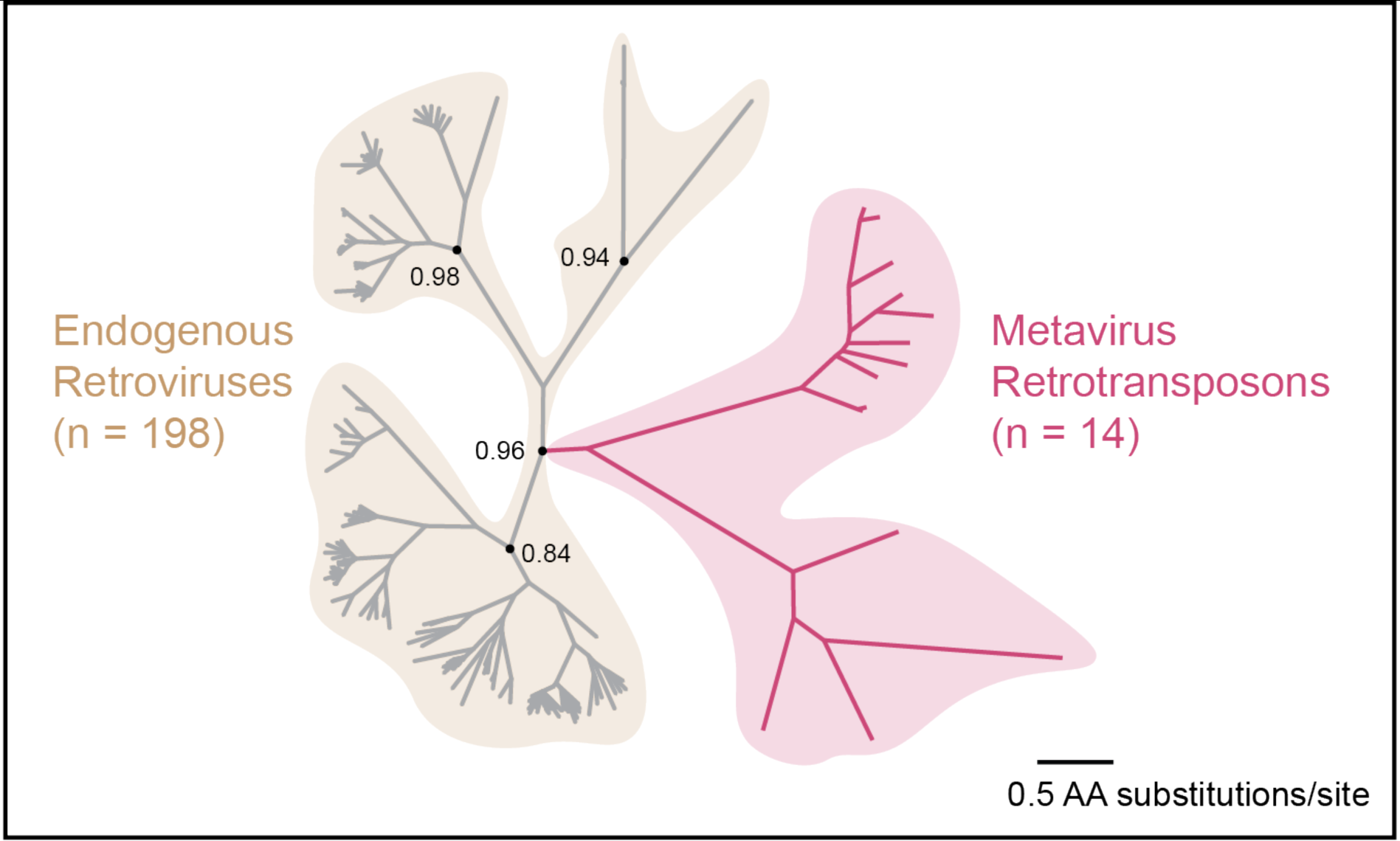
Full-length capsid-like ORFs in the human genome. The human genome contains metaviral capsid ORFs that are divergent and present in low copy number. We generated a maximum-likelihood phylogenetic tree of 212 full-length capsid-like ORFs in the human genome. Hidden Markov Model profile searches identified capsid from three classes of endogenous retroviruses (grey branches), and metavirus retrotransposons (red branches). Maximum likelihood-based support values at selected nodes were calculated in FastTree2.

We found a notable difference in phylogenetic branching patterns between endogenous retroviruses and *Metaviridae*. The majority of endogenous retroviral ORFs cluster with similar sequences related by short branches. This branching pattern is most consistent with differences that arose during recent autonomous replication (Mager and Stoye 2015) but cannot exclude the possibility that some ERV-derived capsid sequences have been recently domesticated; the relative recent replication of ERVs makes it more difficult to distinguish insertions from domesticated copies. In contrast, the single clade representing capsid-like sequences from *Metaviridae* has significantly fewer members, most of which are separated by long branches. This finding is consistent with previous analyses, which showed that *Metaviridae* ceased active transposition in an ancient ancestor of modern mammals (Jürgen Brandt et al. 2005; Blikstad et al. 2008; Mager and Stoye 2015). Thus, any remaining metaviral capsid-like sequences are much more likely to be the result of host domestication (J. Brandt et al. 2005; Kaneko-Ishino and Ishino 2015). Indeed, we found no adjacent long-terminal repeats flanking genomic sequences of metaviral capsids genes using LTRHarvest(Ellinghaus et al. 2008), confirming that these sequences do not represent retrotransposons capable of autonomous transposition. For this reason, we focus the remainder of our study on the 24 intact metaviral-derived capsid-like sequences that remain in the human genome (and one additional pseudogene, a pseudogenized copy of *PNMA6EF* with internal frameshift mutations that are not conserved between human and macaque). Our phylogenetic tree (**Figure 1**) only contains 14 of these 24 *Metaviral* sequences because we excluded duplicates and partial capsid genes. However, we included both full-length and partial capsid-like sequences in subsequent analyses because previous studies have shown that even domesticated metaviral genes encoding a truncated capsid domain can still be functional (Campillos et al. 2006).

Our analyses confirm that previous catalogs of intact human domesticated metaviral capsid genes (Campillos et al. 2006; Kokošar and Kordiš 2013) were complete. These 24 human metaviral capsid-like sequences correspond to previously identified domesticated capsid genes (Campillos et al. 2006; Kokošar and Kordiš 2013), including *ARC*, and members of the *PNMA* and *SIRH/RTL* gene families. The only exception is ASPRV1 (or SASPase) which a previous study identified as containing capsid-like sequence (Campillos et al. 2006), whereas a subsequent study suggested it only has protease but not capsid domains (Kokošar and Kordiš 2013). Consistent with the latter study, our analysis (both HMM analyses and AlphaFold predictions) suggest that ASPRV1 has a protease domains as previously reported but does not have a capsid domain. Previous studies of the *PNMA* gene family also identified four additional members of the family that did not appear among the capsid-like sequences we identified: *CCDC8, PNMA8A, PNMA8B and PNMA8C* (Schüller et al. 2005; Pang et al. 2018). We independently identified these genes via a capsid-adjacent domain (see below) and included these genes in all subsequent analyses because of their close evolutionary ties to the capsid domain in the PNMA family. Multiple gene nomenclatures have been adopted by different studies over the years; to avoid confusion, we list all alternative gene names to help cross-reference previous publications with our results (**Table S2**).

### Domesticated metaviral capsid genes experienced distinct evolutionary retention fates

The 24 human metaviral capsid-encoding genes were domesticated in an ancient mammalian ancestor (Kaneko-Ishino and Ishino 2015) and have subsequently been conserved in at least a few mammalian species (J. Brandt et al. 2005; Naruse et al. 2014; Irie et al. 2015; Irie et al. 2021). We investigated their retention outcomes in a deeper sampling of representative placental mammalian genomes to identify any lineage-specific changes in their retention and investigate the evolutionary constraints and origins of both the capsid and non-capsid domains encoded by these domesticated genes.

We performed searches of 17 additional representative mammalian genomes (14 placental mammals, two marsupials, and one monotreme) using sensitive HMM searches as well as iterative blast searches with both nucleotide and protein queries. We assigned the resulting sequences to orthologous groups using phylogenies and/or sequence similarity. We generated in-frame nucleotide alignments for each group (Methods, see Supplementary Materials for alignment files) and looked for inactivating frameshifts and/or premature stop codons relative to the annotated human ORF. In all cases, the full-length human annotation is well-supported by conservation in distantly related placental mammal genomes (**Figure S2-S5**).

Our phylogenetic analyses (**Figure 2, Figure S2-S5**) confirm previous estimates of the age of these genes (Edwards et al. 2008; Iwasaki et al. 2013; Kokošar and Kordiš 2013; Kaneko-Ishino and Ishino 2015; Pastuzyn et al. 2018a). Most domesticated metaviral capsid genes are at least ∼100 million years (My) old, with orthologs in diverse placental mammals. *PEG10* is slightly older (at least ∼160 My), with additional marsupial paralogs that have been previously described (Suzuki et al. 2007; Ono et al. 2011), whereas *ARC* is much older (at least ∼350 My), with orthologs found across all tetrapods including birds, reptiles, and amphibians (**Figure 2**, **Figure S2**).

**Figure 2.**
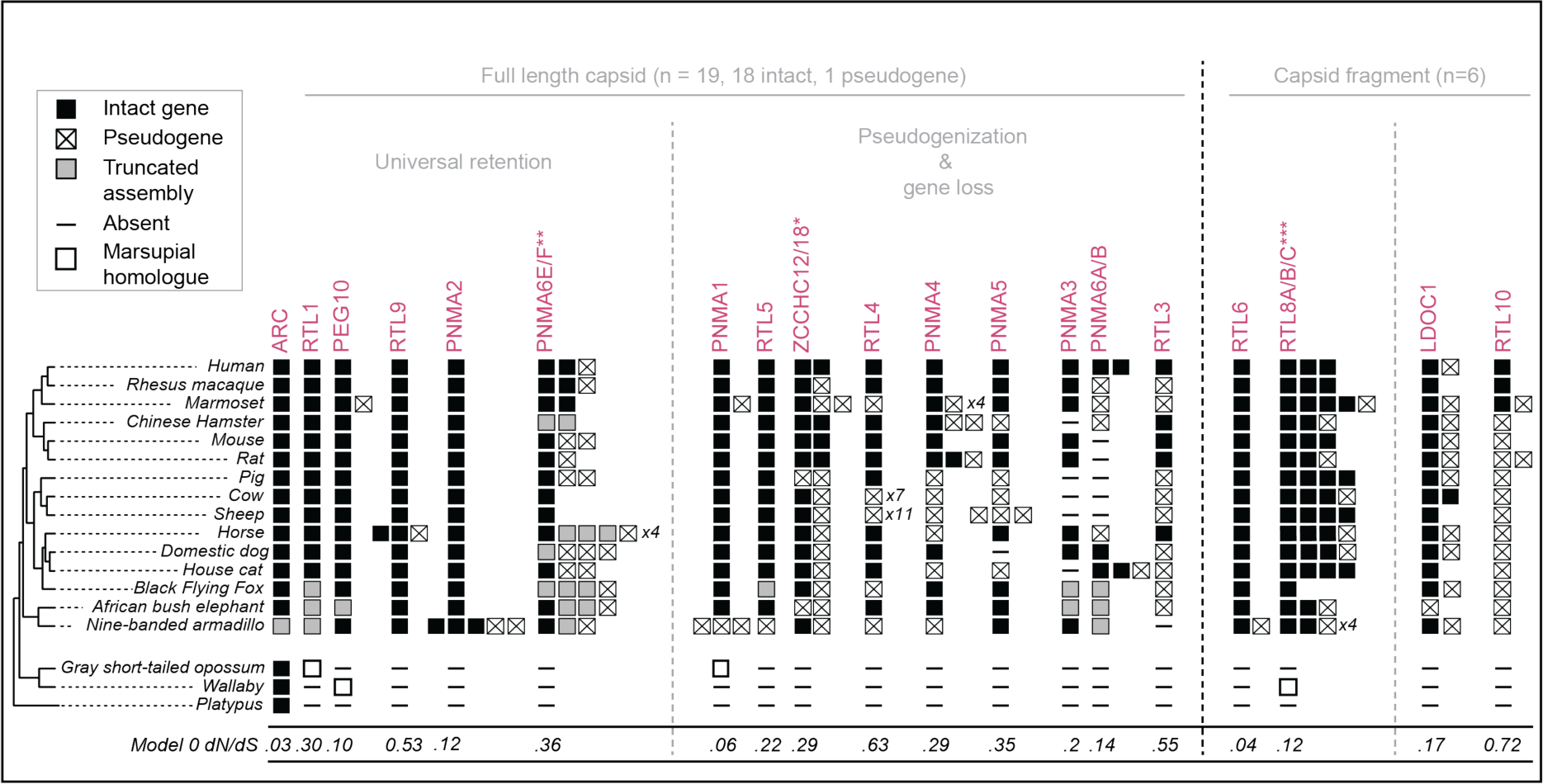
Metaviral-derived genes show distinct evolutionary trajectories across placental mammals. Some full-length metaviral-derived capsid genes are universally retained across placental mammals, whereas others have experienced lineage-specific loss, pseudogenization or duplication events. Filled squares represent intact genes and squares containing a cross represent sequences with obvious inactivating mutations (frameshifts and/or premature stops). Gray boxes represent sequences that are truncated due to genome assembly gaps, and ‘-‘ symbols represent cases where we could find no matching sequence at all. A known species tree is shown on the left and was obtained by pruning whole-genome trees available via the UCSC genome browser. Open boxes represent marsupial sequences identified in our vertebrate genome scan and are aligned beneath their top BLAST hit, which are not necessarily orthologs. Marsupial domesticated genes have been comprehensively addressed elsewhere (Ono et al. 2011; Iwasaki et al. 2013).

However, the repertoire of domesticated genes has not remained static across placental mammals afterautonomous replication ceased. We found that four orthologous groups in our trees (*PNMA6E/F, ZCCHC12/18, PNMA6A/B*, and *RTL8A/B/C*; **Figure 2, Figure S3-S5**) contain more than one human gene (with orthologs present in other species), indicating that more recent lineage-specific duplication events continue to shape some branches of the gene family. For example, *PNMA6E* and *PNMA6F* appear to be the result of a duplication in the common ancestor of primates (**Figure S5**). Primates are not unusual in this regard; many other mammal lineages also appear to have undergone independent duplications in their *PNMA6E/F* family (**Figure S5**). The independent, lineage-specific *PNMA6E/F* duplications could instead be the result of recurrent, lineage-specific gene conversion events, reminiscent of the evolutionary dynamics we previously described in histone variant genes (Molaro et al. 2018) and antiviral IFIT genes (Daugherty et al. 2016). After accounting for lineage-specific duplications, the 24 human domesticated metaviral capsid-like genes group into 19 clades, which likely represents the number of independently domesticated capsid genes in the common ancestor of all placental mammals.

Of these 19 clades, eight are universally retained as at least single intact genes across all placental mammals queried, suggesting important biological function common to all mammals (**Figure 2)**. These eight universally genes contain full-length as well as C-terminally truncated capsid-like regions, confirming that a full-length capsid-like domain is not necessary for the new host-related function of some of these genes (Campodonico et al. 2023). In contrast to the eight universally retained genes, we found that 11 domesticated capsid genes present in the common ancestor of placental mammals have undergone gene loss or pseudogenization in one or more mammalian lineages, suggesting that their function is not universally required in mammals or is redundant with other genes (**Figure 2)**. For example, *RTL10* was present in the common ancestor of placental mammals but is retained as an intact ORF in only the three out of eighteen mammalian genomes we surveyed. Deeper examination in additional species **(Figure S6)** reveals that *RTL10* is also retained under purifying selection (see below) as an intact gene in at least two other diverse mammalian linages: basal glires and cetaceans, where it appears to be subject to purifying selection (below). Thus, even though *RTL10* has been repeatedly lost during mammalian evolution, its retention in some species is nevertheless the result of functional conservation.

Our findings come with caveats associated with genome assembly quality. Despite significant advances since the last compendium of domesticated genes was created, most of these genome assemblies are still incomplete, leading to cases where an assembly gap results in a truncated sequence (*e.g.,* armadillo *ARC*) that is unlikely to be real. This problem is especially compounded in the case of recent duplicates, which are likely undercounted in *de novo* assemblies that are more likely to contain assembly gaps (**Figure 2**). Finally, genome assemblies contain small numbers of sequencing errors, so a minority of the apparent pseudogenes could represent intact genes with sequencing errors. Given these caveats, where possible, we used additional sequences from sister lineages to gain more confidence that inactivating mutations are not simply genome assembly or sequencing errors.

### Domesticated metaviral capsid genes are retained under purifying and positive selection

The cadence and locations at which amino acid changes accumulate during evolution can also provide important clues about a protein’s cellular function. We investigated the selective pressures on metaviral genes following their domestication (**Figure 2**, **Table 1**), estimating the ratio of non-synonymous (dN, amino acid altering) to synonymous (dS, amino acid preserving) nucleotide substitutions using the codeml algorithm from the PAML suite (Yang 2007). Most protein-coding genes exhibit low dN/dS (<< 1), indicating purifying selection, where non-synonymous (amino acid altering) mutations are less likely to be tolerated than synonymous (amino acid-preserving) mutations. dN/dS close to 1 indicates neutral evolution (no protein-coding constraint), and dN/dS >1 indicates positive selection (adaptive evolution favors amino acid changes).

**Table 1:**
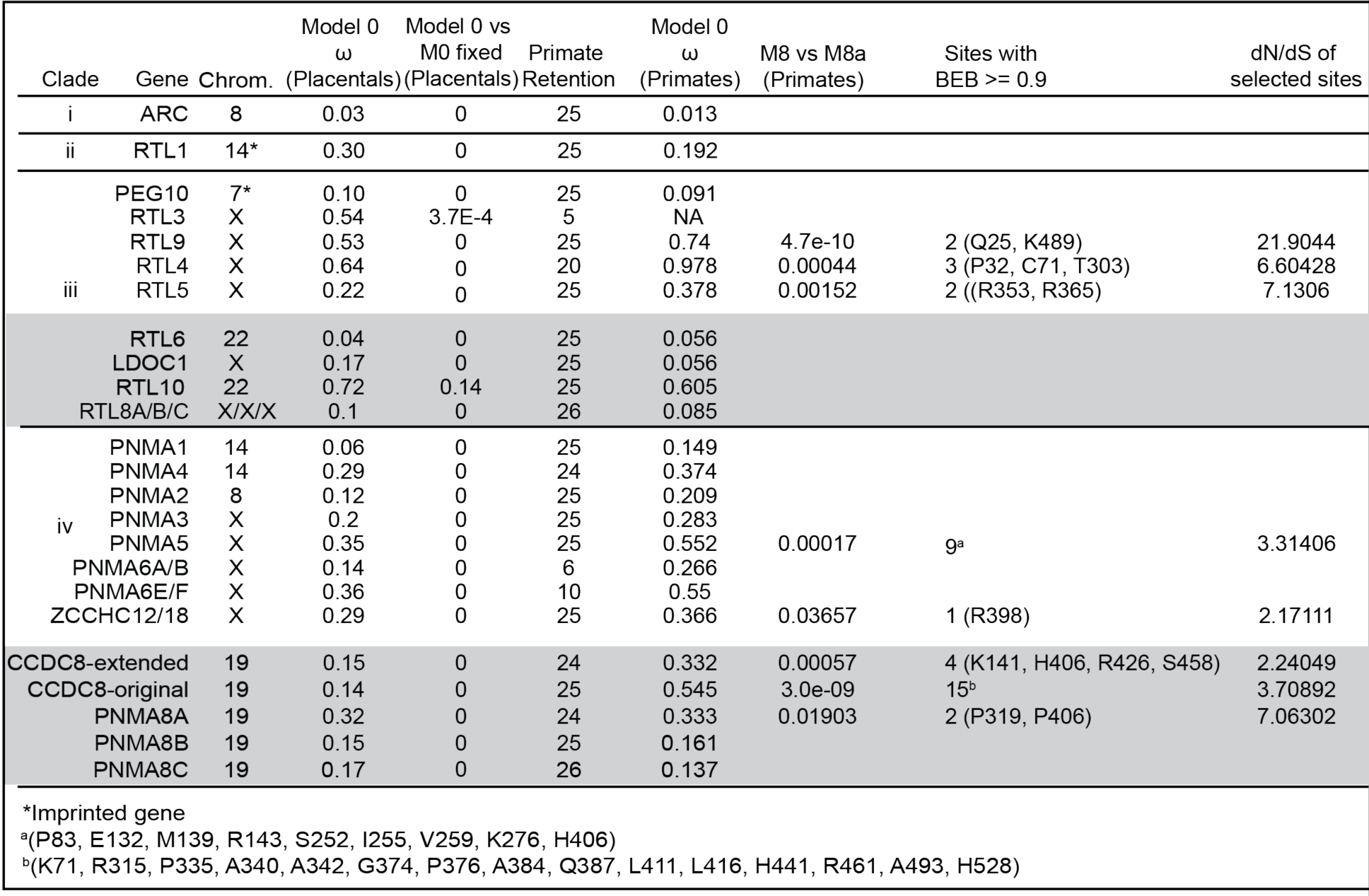
Evolutionary rates of domesticated metaviral genes across placental mammals and primates. Phylogenies for each gene were generated using PHYML with GTR substitution model. Alignments and trees were used as input to the codeml algorithm in PAML. To assess rapid evolution, we analyzed genes from primates alone. Residues identified in primates as evolving under rapid evolution are shown in parentheses, with position in the human gene.

We first estimated the overall dN/dS of alignments of intact ORFs from across placental mammals (n=18 species), using PAML’s model 0 that makes the assumption of uniform selective pressures across all sites and all lineages. We compared the likelihood of this observed dN/dS compared to a model 0 with fixed dN/dS of 1, allowing us to test whether the observed dN/dS model was a better fit than neutral evolution. We find that most *Metaviridae*-origin capsid genes evolve under overall purifying selection, with *ARC* showing some of the strongest signatures of purifying selection (dN/dS=0.03). In contrast, for *RTL10*, we were barely able to rule out the null hypothesis of neutral evolution (dN/dS =1) using our initial sampling (n=3 intact primate genes). However, our deeper alignments for *RTL10* provided greater statistical power, revealing evidence of purifying selection in basal glires (rodents/lagomorphs) and in cetaceans, despite pseudogenization in other lineages (**Figure S6)**. It remains possible that in some lineages, *RTL10* could have functions that are unrelated to its protein-coding capacity, for example as a non-coding RNA. Indeed, preliminary data from the International Mouse Phenotyping Consortium (see below) indicates that mouse knockouts of *RTL10* have both behavioral and morphological phenotypes, despite apparent frameshifting mutations in the capsid-encoding region of the mouse gene (**Figure S6)**. Nevertheless, *RTL10* represents an unusual case; for most domesticated genes, there is unambiguous evidence of their retention under purifying selection. Overall, our evolutionary analyses indicate that most domesticated metaviral capsid genes have been retained under strong selective pressures during the last 110 million years of placental mammal evolution, suggesting that they each play important, non-redundant roles in mammalian biology.

Despite an overall signature of purifying selection, domesticated capsid genes might still be subject to positive selection (dN/dS >1) at a subset of codons as seen for host immunity related genes such as *Fv1* (Yap et al. 2014; Young et al. 2018). To scrutinize each codon, we collected sequences from the simian primate lineage, because this taxonomic group provides good statistical power to detect site-specific positive selection (McBee et al. 2015). We analyzed these alignments using PAML’s maximum likelihood tests that ask whether evolutionary models that allow a subset of sites with dN/dS>1 (NSsites model 2 or 8) are a better fit to the observed sequence data than matched models that only allow sites under neutral and purifying selection (dN/dS of 0-1) (NSsites models 1, 7, or 8a). These tests reveal that eight ancient metaviral genes contain a subset of sites evolving under positive selection in primates (**Table 1**), suggesting ongoing involvement in evolutionary arms races. Our findings of positive selection are intriguing given previous studies that implicate three of the positively selected genes (*CCDC8, RTL5, RTL9*) in immune-related functions (Wei et al. 2015; Irie et al. 2022; Ishino et al. 2023), implying that host-pathogen genetic conflict may be driving the rapid evolution of at least some of these genes.

### Domesticated capsid genes are structurally diverse

Next, we compared the domain architecture of metaviral-derived human genes to those of their metaviral progenitors, taking advantage of significant recent advances in protein structure prediction and structural homology search algorithms (Holm 2020; Jumper et al. 2021; Kempen et al. 2022). To identify metaviral relatives, we queried non-mammalian vertebrate lineages where *Metaviridae* are still actively retrotransposing, using six-frame translations of the chicken (*Gallus gallus),* alligator (*Alligator mississippiensis),* painted turtle (*Chrysemys picta bellii)*, anole lizard (*Anolis carolensis*), African clawed frog (*Xenopus laevis*), and coelacanth (*Latimeria chalumnae*) genomes. We aligned these metaviral capsid sequences with the mammalian capsid-like genes and generated a maximum-likelihood phylogenetic tree of the resulting 781-sequence alignment (**Figure 3a**).

**Figure 3.**
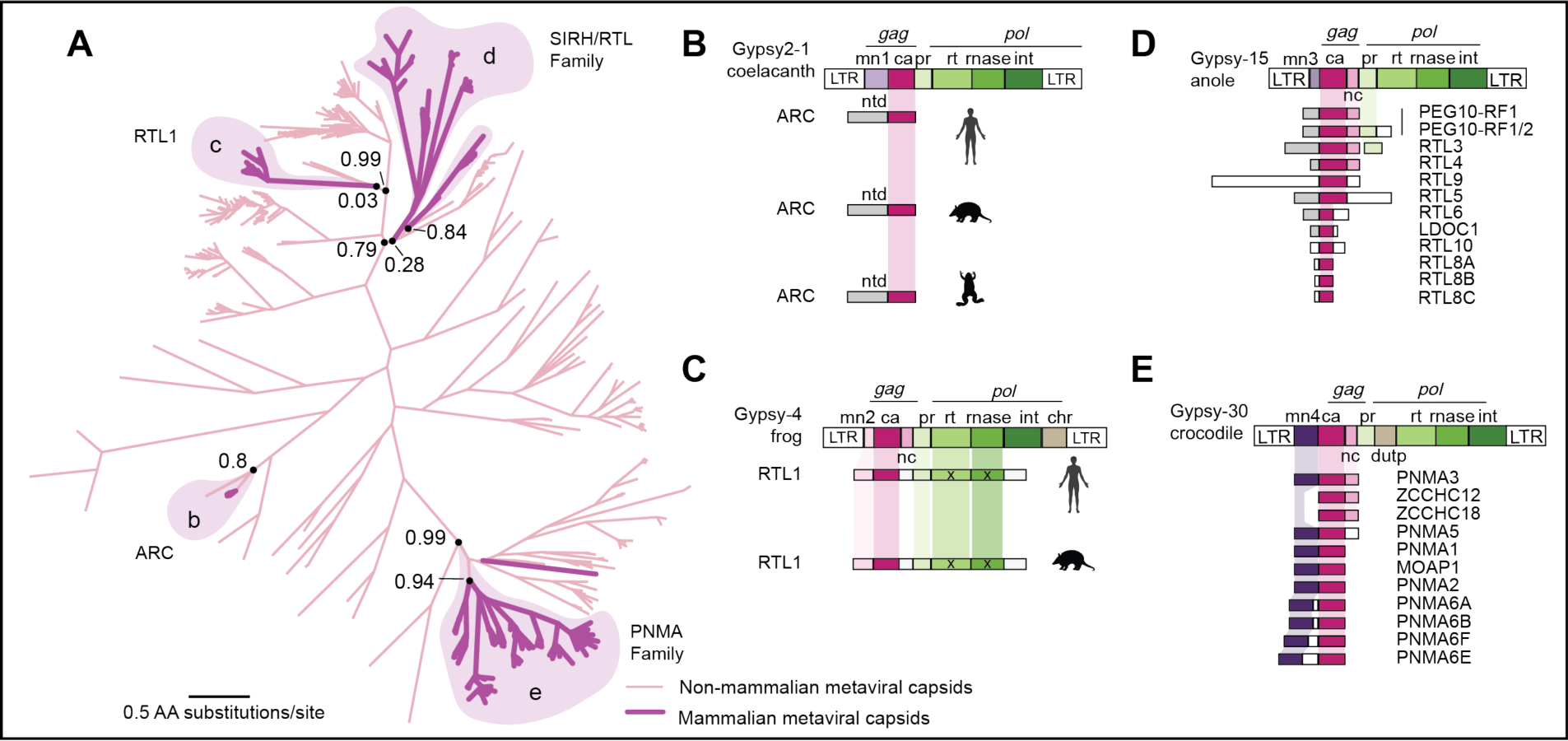
Four independent metavirus domestication events include structurally distinct N-terminal domains. **A.** A maximum likelihood phylogenetic tree of 781 capsid sequences from 24 vertebrate species. The tree includes 119 domesticated metaviral capsid genes found in diverse mammals (dark purple, highlighted) and 662 metaviral capsid-like ORFs (light pink) from selected non-mammalian vertebrate species (chicken (n=1), alligator (n=13), painted turtle (n=32), anole lizard (n=310), African clawed frog (n=291) and coelacanth (n=15). Maximum likelihood-based support values at selected nodes were calculated in FastTree2. **B-E.** Domain architecture (not to scale) of human *Metaviridae*-derived capsid genes and the closest available consensus metaviral sequence from Repbase, organized according to major clades in the tree shown in panel A. Colored boxes indicate domains within each open reading frame identified by HMM profile searches, structural prediction, and structural homology searches.

Consistent with previous studies (Campillos et al. 2006; Kokošar and Kordiš 2013), our phylogeny separates the domesticated metaviral genes into at least 4 separate clades, each containing (likely active) metaviral outgroups from non-mammalian genomes; in our subsequent analysis, we use these modern, active metaviral sequences as “ancestral proxies” for the ancestral metavirus that was originally domesticated. Two clades are represented by single mammalian genes (*ARC* and *RTL1*), whereas two other clades (*PNMA* and *SIRH/RTL*) have expanded to contain many mammalian members, demonstrating that gene duplications (rather than independent retrotransposition events) probably led to the expansion of each gene family (Campillos et al. 2006; Kokošar and Kordiš 2013).

Many of the domesticated metaviral capsid genes include additional domains, which sometimes include additional metaviral-derived regions, as well as protein segments with no recognizable viral homology (Campillos et al. 2006; Kokošar and Kordiš 2013). To characterize these domains, we first focused on annotating non-capsid domains of ‘ancestral proxy’ metaviruses found in our genome scan and consensus sequences from the database Repbase (Bao et al. 2015) using LTR-Harvest, HMMER, PFAM (now part of Interpro), Alphafold, Foldseek, and DALI (Ellinghaus et al. 2008; Eddy 2011; El-Gebali et al. 2019; Holm 2020; Jumper et al. 2021; Kempen et al. 2022). We then compared the resulting patterns to those observed in the mammalian domesticated metaviral genes. For the domesticated genes, we also used these additional non-capsid HMMs to search 6-frame translations of 16kb of flanking genomic sequence on each side of the capsid region, to rule out the possibility of any gene mis-annotation or subsequent insertions having split and separated the original metaviral domestication. Several of our annotations agree with previously published observations (Campillos et al. 2006; Kokošar and Kordiš 2013). For completeness, we summarize previous observations as well as discuss our findings below.

First, while many metaviral gag-derived genes encode a complete capsid domain, several in the *RTL/SIRH* family encode a truncated capsid domain that has only the N-terminal lobe (*e.g.,* LDOC1). These truncations are seen across mammalian orthologs of each gene, indicating that they occurred soon after gene birth **(Figure 3, Figure S7)**. Notably, the AlphaFold predictions indicate that proteins encoded by these ‘half-capsid’ genes have absolutely conserved the structural fold of the N-lobe from the ancestral metavirus, whereas adjacent domains have diversified, and are typically predicted to be a mix of single alpha helices and disordered regions of varying lengths (schematicized in **Figure 3**, **Figure S8)**. A recent report suggests the N-lobe encoding RTL8 regulates the function of the full-length capsid PEG10 (Campodonico et al. 2023). Other truncated capsids may perform analogous regulatory functions of full-length capsid proteins.

Second, several RTL/SIRH family genes, as well as RTL1, encode full-length metaviral capsid domains fused to long N- or C-terminal unstructured regions (white boxes, **Figure 3D**). Our in-depth structural analysis finds no additional recognizable domains in these regions and no evidence that these regions are metaviral-derived. Although these regions are likely unstructured, they are nevertheless well conserved across placental mammalian orthologs (**Figure 3b-d, Figure S4**), indicating their functional importance.

Third, PEG10 encodes a protease domain after a -1 ribosomal frameshift that is nearly universally conserved in placental mammals; ∼30% of translating ribosomes undergo the frameshift, resulting two different protein products: a shorter majority product containing only the *gag*-related region, and a longer minority product that also contains a protease domain (Shigemoto et al. 2001; Clark et al. 2007; Shiura et al. 2021). RTL1 proteins encode protease, and (likely catalytically inactive but structurally conserved) reverse transcriptase and Ribonuclease H (RNase H) domains (Lynch and Tristem 2003).

Like *PEG10*, a programmed frameshift and downstream protease domain have also been suggested for the *RTL3* gene (Jürgen Brandt et al. 2005; J. Brandt et al. 2005), although the functional relevance or evolutionarily history has not been well-defined (*e.g.,* it is not noted in the RefSeq annotations). We used numerous additional placental mammal genomes to investigate *RTL3*’s putative programmed frameshift (**Figure S6**). Although *RTL3* is clearly a pseudogene in several major mammalian clades (*Carnivora*, *Bovidae*), both protease and capsid-containing ORFs remain intact in many other lineages. In these instances, the protease domain is very well conserved and shows much stronger evidence of amino acid constraint (overall dN/dS=0.11) than the capsid-containing ORF (overall dN/dS=0.67; dN/dS of capsid domain alone=0.55) (**Figure S6**). In the mouse *Rtl3* ortholog, a simple -1 frameshift would result in a fusion protein that contains both capsid and protease domains. However, in human *RTL3*, a -1 frameshift would not be sufficient to produce a fusion protein, because the stop-free regions containing the two domains do not overlap and stop codons exist in all three reading frames in the intervening regions. Furthermore, in many other simian primate species, one or both domains of RTL3 clearly acquired inactivating mutations in ancestral species (**Figure S6**). The most likely explanation is that *RTL3* encodes as a capsid-protease fusion protein in mouse and many other mammalian lineages, but not in human and other simian primates. Instead primate *RTL3* encodes independent capsid and protease proteins. Similar retention of programmed frameshifts have also been reported for *PNMA3* and *PNMA5* (Wills et al. 2006). However, sequence following the frameshift is not as well conserved as that in *PEG10* or *RTL3* and we found no evidence for functional domains downstream of these putative frameshift sites; thus, their functional relevance remains unclear.

### Diverse N-terminal structures in domesticated genes derived from metaviral ancestors

The most surprising finding from our reannotation emerged from the analysis of the four ‘ancestral proxy’ metaviruses, each of which has a distinct domain architecture **(Figure 3b-e)**. Of the four, only the ARC ancestral Metavirus did not contain a nucleocapsid (NC) domain immediately downstream of the capsid domain. In contrast, a recognizable NC is present in the other three ancestral metaviruses and has been maintained in some but not all domesticated genes in these three clades. NC domains bind viral RNAs to help package them into capsid (Muriaux and Darlix 2010). Their presence in some domesticated genes suggests that encoded proteins that are more likely to have RNA-binding capabilities, as has been demonstrated for PEG10 (Abed et al. 2019; Segel et al. 2021). However, a NC domain is not apparently necessary for RNA packaging; ARC packages RNA even though it lacks a canonical nucleocapsid domain (Pastuzyn et al. 2018). The four ‘ancestral’ metaviruses also differ in some other idiosyncratic ways; a chromodomain is found only in the *RTL1* ancestor, and dUTPase only in the *PNMA* family ancestor. However, neither the chromodomain nor the dUTPase was retained in domesticated genes, so we do not study these domains further.

We found that the N-terminal domains of each of these four ancestral metaviruses were completely dissimilar to each other (**Table S3**), revealing a previously underappreciated degree of domain complexity in a poorly studied region of the *gag* gene of metaviruses and retrotransposons in general. Previous analyses have described the N-terminal region of metaviral ORFs as matrix domains, by analogy to retroviral matrix domains that function to facilitate membrane association and virion formation (Campillos et al. 2006; Kokošar and Kordiš 2013; Pastuzyn et al. 2018). However, this domain assignment has been based more on analogy than actual homology. Typically, retroviral matrix domains fold into globular core composed of four alpha helices with a conserved basic surface patch that facilitates membrane interaction (Murray et al. 2005; Hamard-Peron and Muriaux 2011). We explored the domain architecture of both the four ‘ancestral proxy’ metaviruses and derived domesticated genes using a combination of BLAST, AlphaFold, DALI and Foldseek. We were surprised to find no evidence of a canonical retroviral matrix domain in any of the four metaviruses. Indeed, in our survey of actively circulating metaviral retrotransposons, we find the N-terminal domains (*i.e.,* upstream of the capsid) to be surprisingly variable in length, and typically predicted by AlphaFold to be a mix of single long alpha helices and disordered regions (**Figure S8**).

Given their distinct evolutionary origins and domain architecture, we examined each of the four clades of domesticated genes in more detail to infer functions. We first examined the ∼350 My-old ARC gene, analyzing the mammalian orthologs described above (**Figure 2**), as well as a more distant frog ortholog. Notably, ARC’s entire gene architecture is conserved across species, not only the capsid domain. However, upon comparison of ARC with ancestral’ ‘proxy’ metaviral sequences from the coelacanth, we found homology between capsid domains but not in the N-terminal region, which in ARC is a ∼200 amino acids long coiled-coil domain (using reciprocal HMM searches; queries were N-terminal HMMs built either from 11 coelacanth metaviral sequences, or from vertebrate ARC orthologs) **(Figure 3b)**. Furthermore, we failed to find matches to any metaviral retrotransposons in the database Repbase using the ARC N-terminal region HMM. Additional efforts using Hhpred (Zimmermann et al. 2018), DALI(Holm 2020), and FoldSeek (Kempen et al. 2022) also failed to clarify the evolutionary origin of this 200 amino acid domain of ARC. We cannot rule out the possibility that this unusual N-terminal domain derived from a metaviral lineage that has not yet been sequenced, especially since the domestication event is so old. Alternatively, it remains possible that the domain was acquired at or following domestication; we note however that no homologous matches were found in any animal genomes. Thus, while the capsid domain is unambiguously of metaviral origin, the origin of the ARC N-terminus remains mysterious and should be a focus given the importance of this domain in ARC function (Eriksen et al. 2021).

The N-terminal regions of RTL1 and the RTL/SIRH family (and their closest metaviral relatives) encode single alpha helices and disordered regions ranging from 25 residues (RTL8) to 1169 residues (RTL9) with no clear domain architecture. These N-termini are conserved across placental orthologs but are not similar between the genes in this family, suggesting they are functionally important domains, despite our inability to predict their function through structural homology (**Figure S8**). This reveals that the N-terminus of the *RTL/SIRH* family has diverged widely since domestication, while the capsid core has retained nearly perfect homology to a metaviral capsid domain. This contrasts starkly with the fate of the N-terminus in the *PNMA* family, which contains a highly conserved N-terminus, which we discuss in more detail in the next section.

### A novel RNA-binding domain in the PNMA family

Structural modeling and similarity searches of the ∼150 amino acid N-terminal region in the *PNMA* clade and its closest metaviral relative revealed an ∼100 amino acid RNA-binding domain (RBD) with mixed alpha-helical and beta sheet topology that was previously unrecognized in any metaviral sequences or in the domesticated *PNMA* gene family (**Figure 3e**, **Figure 4**). Using AlphaFold’s structural prediction for the N-terminal domain from human PNMA1 as a query in DALI and Foldseek structural homology searches, we found high-scoring matches to a widely distributed RNA-binding domain found in many proteins, including an RNA recognition motif of the p65 subunit of the *Tetrahymena thermophila* telomerase complex (PDB: 7LMA)(He et al. 2021) and human MARF1 (meiosis regulator and mRNA stability factor 1, PDB: 2DGX, no structure-associated publication) (**Figure 4B**). Due to several missing residues in the Gypsy-30 consensus sequence, we instead used the closely related and complete *Danio rerio* Gypsy-12 consensus sequence in the Repbase database (Bao et al. 2015) for structural predictions (**Figure S10A**). This prediction demonstrates clear structural homology with both PNMA1 and the same RNA-binding domains (henceforth RBD), again demonstrating that this domain was present in the ancestral metavirus before domestication in mammals.

**Figure 4.**
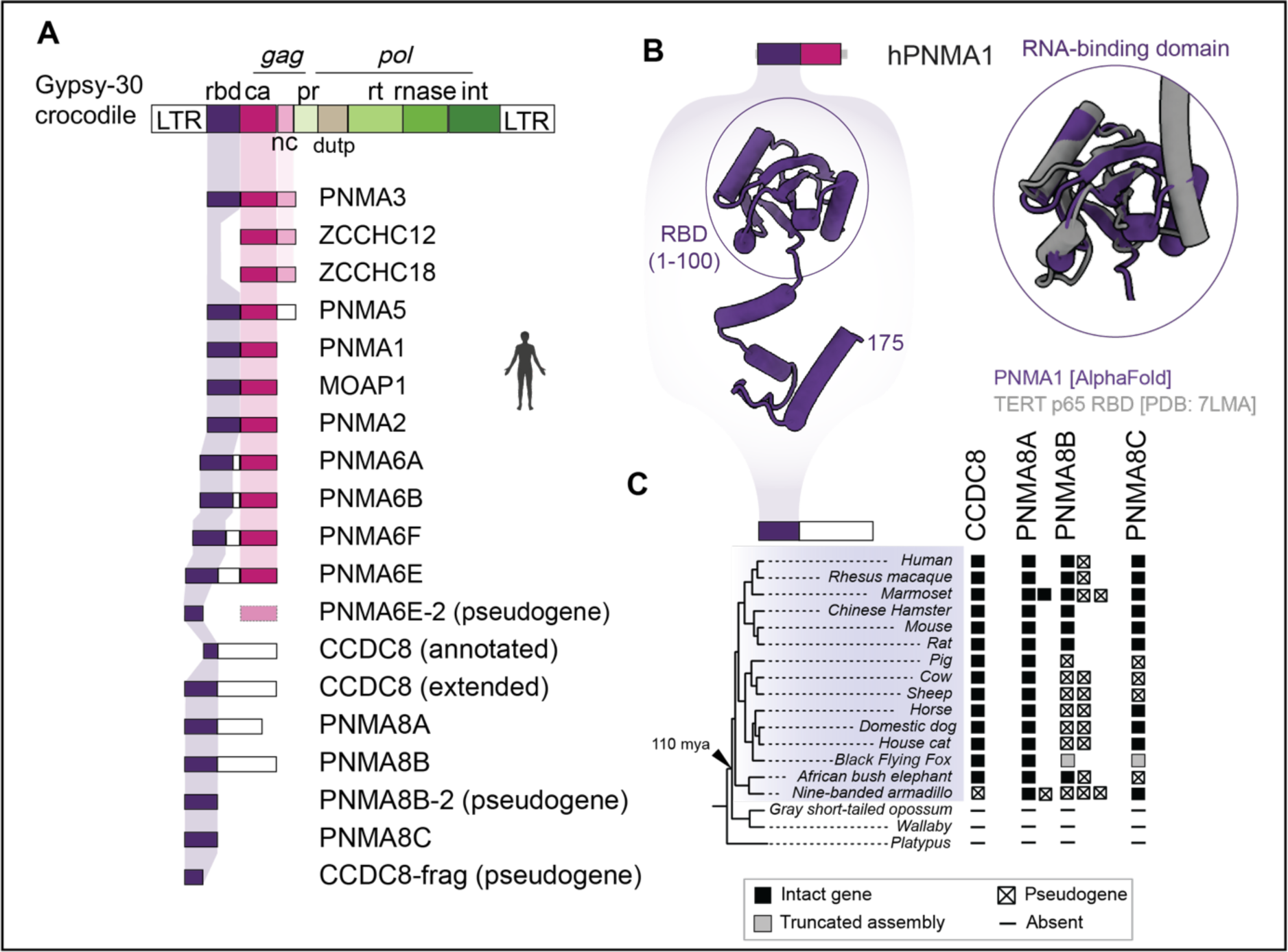
A conserved RNA-binding domain in the PNMA family and related metaviruses. **A.** A metaviral-derived RNA-binding domain is present N-terminal to the capsid domain in nine domesticated genes and in isolation in four additional genes (human gene architectures shown). **B.** AlphaFold structural prediction of PNMA1 (blue), shown alone (left) as well as superimposed on an experimentally determined structure (gray, right) of the telomerase p65 protein’s RNA binding domain (PDB: 7LMA) (RMSD between 41 C*α* atoms is 1.2 å and 6.4 Å across all 73 equivalently positioned C*α* atoms) **C.** Retention of RBD-only genes in placental mammals. Filled squares represent intact genes and squares containing a cross represent sequences with obvious inactivating mutations (frameshifts and/or premature stops). Gray boxes represent sequences that are truncated due to genome assembly gaps, and ‘-‘ symbols represent cases where we could find no matching sequence at all. A known species tree is shown on the left and was obtained by pruning whole-genome trees available via the UCSC genome browser

We wished to ensure that we could identify all instances of human proteins that contain this novel RBD domain. Therefore, we generated an HMM and performed several sensitive database searches. Searching the 6-frame human genome translation, we found almost all *PNMA* family members, except *ZCCHC12* and *ZCCHC18,* encode the RBD domain. Nine of the human RBD-containing genes also contain a capsid domain, but four do not. These four non-capsid genes (*CCDC8, PNMA8A, PNMA8B and PNMA8C*) represent the four metaviral derived genes that we had originally missed based on our capsid-focused HMM search strategy. Thus, the only domain that links these four genes to the rest of the members of the domesticated PNMA gene family is the novel RBD that we have identified. Three of these non-capsid genes encode proteins in which the RBD is fused to unstructured C-terminal regions. We also performed the same cross-species analysis as before to investigate the evolutionary retention of the RBD. We found that all *PNMA* family members, including capsid-encoding and capsid-lacking genes, show significant conservation of the RBD across placental mammals **(Figure 4**, **Table 1)**.

Our identification of the capsid domain also allowed to us to correct a major mis-annotation of one of the *PNMA* gene family members. While most of the HMM matches spanned the full length of the RBD, our initial survey found only a partial match to the RBD domain in the annotated *CCDC8* protein, suggesting it contained an apparently truncated form of the RBD domain. This truncation seemed surprising, so we explored *CCDC8* genomic sequences more closely. We found strong evidence that the correct initiation codon for *CCDC8* translation is at a non-canonical upstream CTG start codon, encoding a 608 amino acid protein containing a full-length RBD match **(Figure S9)** instead of the currently annotated 538-residue protein. Thus, identifying the novel RBD allowed us to correct the annotation of the *CCDC8* gene.

The *PNMA* gene architecture demonstrates that a metavirus containing an RBD-capsid-NC domain architecture was active in an ancient mammalian ancestor. By aligning RBD-like sequences and generating a maximum likelihood phylogeny, we conclude that the RBD-like domestication of *PNMA* genes most likely occurred only once and was spread by subsequent duplication **(Figure S10)**, in agreement with inferences based on the capsid domain **(Figure 3)**(Kokošar and Kordiš 2013). To understand how prevalent similar architectures might be in extant metaviruses, we used the same *PNMA* family N-terminal HMM to better understand the phylogenetic distribution and origin of this novel RBD domain. We searched the Repbase database, where we identified 10 consensus *Metaviridae* containing the RBD. We also searched six-frame translations of genomes in which metaviruses are still active: alligator (*Alligator mississippiensis),* painted turtle (*Chrysemys picta bellii)*, anole lizard (*Anolis carolensis*), African clawed frog (*Xenopus laevis*), coelacanth (*Latimeria chalumnae*), zebrafish (*Danio rerio*), and fugu (*Takifugu rubripes*), yielding 217 RBD-like ORFs, including representatives from all genomes searched. These results indicate that, though rare, actively retrotransposing metaviruses containing a RBD-capsid-NC architecture still exist in reptile, amphibian, and fish lineages. The persistence of this RBD as a metaviral domain and the domain’s widespread conservation in domesticated PNMA-family genes highlight the importance of this domain, though its function remains unknown. Thus, our detailed reanalysis of domesticated genes reveals an unanticipated diversity of domain architectures in metaviruses.

## DISCUSSION

Domestication of genes derived from viruses and transposons can significantly expand the coding potential of host genomes. Our reanalysis of *Metaviridae-*derived capsid-like genes in human and placental mammal genomes reaffirms the broad conclusions of previous studies (Campillos et al. 2006; Kaneko-Ishino 2012; Kokošar and Kordiš 2013; Kaneko-Ishino and Ishino 2015; Pang et al. 2018) and identifies several previously undiscussed points of interest. Like in previous analyses, we find 24 human genes derived from at least four independent germline integrations from ancient, diverse Metaviruses. However, the evolutionary fate of these domesticated genes varies widely following domestication. Two ancient germline integrations of metaviruses created universally conserved single-copy genes (*ARC, RTL1*), whereas two others serially duplicated to become the *SIRH/RTL* and *PNMA* gene families (Campillos et al. 2006; Kaneko-Ishino 2012; Kokošar and Kordiš 2013; Kaneko-Ishino and Ishino 2015). Almost all genes reveal signatures of having been preserved via purifying selection in some mammalian lineages. However, our re-examination reveals only 11 of 24 domesticated genes are universally conserved among placental mammals (8 sets), suggesting these 11 genes perform important functions that have remained largely unchanged in ∼100 My of mammalian evolution. In contrast, other domesticated genes have experienced lineage-specific losses or duplications, suggesting lineage-specific function. For example, several of the non-universally retained domesticated genes are known to function in placenta or the brain, both organs that exhibit huge anatomic and functional diversity among mammals (Cho et al. 2008; Takaji et al. 2009; Irie et al. 2022). Moreover, some of these genes evolve under positive selection, consistent with their involvement in evolutionary arms races and recent reports of immune-related functions (Jiang et al. 2020; Irie et al. 2022; Ishino et al. 2023). However, it is unclear whether immune-related functions can explain all cases of domesticated genes evolving under positive selection.

Domesticated metaviral genes are expressed in diverse tissues. Systematic analysis of gene expression patterns from across 54 adult human tissues (Genotype-Tissue Expression Consortium (GTEx)), and placenta from the Human Protein Atlas and an independent study (Lonsdale et al. 2013; Sjöstedt et al. 2020; Gong et al. 2021) enabled us to compare expression patterns for different domesticated genes (**Figure S11, Table S4**). Some domesticated genes have a widespread expression pattern in all or most human tissues (*e.g., RTL8C, PNMA1*), whereas others are highly tissue-restricted in their expression (*e.g., PNMA6F* is expressed principally in brain, *RTL9* principally in testis, and *RTL1* principally in placenta). We found robust evidence of expression for all but one domesticated gene in at least one human tissue. Only *RTL4* does not exhibit robust expression in any of the tissues sampled, which could be a limitation of tissues sampled by GTEx, or be under the limit of detection for bulk RNA-seq.

What are the functions of these domesticated genes? Although the organismal and cellular functions of most domesticated metaviral genes remain functionally uncharacterized, important inroads into functional characterization have been made for several metaviral genes. *ARC* has been extensively studied for its role in learning and memory (Carmichael and Henley 2018; Epstein and Finkbeiner 2018; Mabb and Ehlers 2018; Newpher et al. 2018; Nikolaienko et al. 2018; Okuno et al. 2018). *ARC* knockout mice exhibit deficits in learning and memory and sleep (Plath et al. 2006; Manago et al. 2016; Suzuki et al. 2020). However, the exact role of the capsid domain remains unclear. In addition, Kaneko-Ishino, Ishino and colleagues have generated mouse knockouts of *PEG10, RTL1, LDOC1, RTL4*) and fluorescent reporter knock-in alleles of *RTL5, RTL6* to study the function of metaviral capsid genes in mice. These in-depth studies reveal that many domesticated genes have profound functional consequences on organismal fitness. For instance, loss of *PEG10* leads to complete early embryonic lethality, *RTL1* loss leads to partial lethality at late fetal/early neonatal stages and behavioral phenotypes in surviving animals, whereas loss of *LDOC1* leads to abnormal placental morphology (Ono et al. 2006; Kagami et al. 2008; Sekita et al. 2008; Naruse et al. 2014; Kitazawa et al. 2017; Kitazawa et al. 2021; Chou et al. 2022). Additional studies suggest other metaviral-derived genes function in innate immunity or cognition (Takaji et al. 2009; Irie et al. 2015; Irie et al. 2021; Chou et al. 2022). Complementing these in-depth studies focused on individual genes, eight of nine knockouts of domesticated genes generated and phenotyped by the International Mouse Phenotyping Consortium reveal significant behavioral and/or physiological phenotypes (**Table S5**). However there is little overlap between the genes knocked out by the IMPC KO and other studies – only *RTL4* has been knocked out by both the IMPC and in an independent study (Irie et al. 2015), and in this single instance the high throughput results and detailed study do not agree. Irie and colleagues were able to identify differences in noradrenaline levels in RTL4 knockout mice that they associate with increased impulsivity, reduced attention, and memory deficits that were not observed in the high throughput analysis.

In addition to their native endogenous functions, aberrant expression of some domesticated metaviral genes can lead to autoimmune disease. For example, the *PNMA* gene family is so named because *PNMA1* encodes a protein associated with the autoimmune disorder paraneoplastic syndrome (PNMA1: paraneoplastic Ma antigen 1). Other PNMA genes may have similar phenotypes, although it is unclear currently whether their autoimmune consequences are related to their endogenous function or irregular expression or regulation (Dalmau et al. 1999; Schüller et al. 2005; Henry 2019; Xu et al. 2023). Little is known about the endogenous cellular function of the *PNMA* family genes. Previous studies have revealed that *PNMA4* (*MOAP1)* likely plays a role in regulating apoptosis, but beyond these studies in cancer cell lines, functional studies remain sparse (Tan et al. 2001; Tan et al. 2005; Fu et al. 2007; Foley et al. 2008; Huang et al. 2012; Law et al. 2015). In summary, mouse knockout studies indicate that domesticated metaviral genes perform important functions; their loss leads to profoundly deleterious outcomes from embryonic lethality to cognitive deficits to organ abnormalities. These studies suggest that the domestication and subsequent proliferation of metavirus-derived genes led to novel non-redundant and essential functions (Naruse et al. 2014; Irie et al. 2015; Irie et al. 2021; Chou et al. 2022). How much of their molecular function are directly the result of their metaviral origins (*e.g.,* capsid mediated transport of RNA by ARC and PEG10) remains an active area of investigation in many labs.

Our study also reveals novel insights into the domain architecture of *Metaviridae*-derived domesticated genes. Previous sequence homology studies showed that adjacent metaviral domains (*e.g.,* protease RT, RNase H) have been occasionally retained alongside the capsid-like region, that unstructured regions with no recognizable homology have been added to metaviral-derived sequences of some genes, and that some domesticated capsid-like genes retain only one of the two capsid lobes (J. Brandt et al. 2005; Campillos et al. 2006; Kokošar and Kordiš 2013). Our analysis using recently developed structural prediction and homology search tools further reveal that each of the four ancient domestications has a distinct N-terminal region, with no recognizable homology to retroviral matrix domains. Furthermore, our analyses reveal that one of the four ancestral viruses and most of its descendant domesticated genes (the *PNMA* family) have an N-terminal region with clear structural homology to RNA-binding domains (RBDs). While most PNMA family members retain both a capsid-like and an RBD-like domain, several have lost the capsid but still contain the RBD, often fused to predicted disordered regions. Cross-species conservation, as well as a knockout mouse for *CCDC8*, indicate that even these RBD-only genes can have important functions. Indeed, RBD retention in the domesticated *PNMA* family genes like *CCDC8* further suggests that RNA interactions are an important feature of their present-day host functions, independent of whether they contain an adjacent capsid-like region. Although the previous focus has been on *Metaviridae*-derived capsid domains, our findings highlight an RNA-binding domain of unknown function within the *Metaviridae gag* domain that has been captured in essential protein-coding genes in placental mammals and even amplified via gene duplication.

The *PNMA*-encoded RNA-binding domain (RBD) is present in some modern *Metaviridae* that continue to actively circulate in amphibians, reptiles, and fish. Thus, our analysis of domesticated genes not only reveals an unprecedented diversity of domain architecture in Metaviridae, but also identifies a previously undescribed RNA-binding domain in vertebrate metaviral retrotransposons that has an unknown role in the metaviral lifecycle. Viral capsid domains are already known to function to package viral RNA genomes, and the C-terminal nucleocapsid is already predicted to bind RNAs. The prevalence of a tripartite RBD-capsid-NC architecture in ancient and active metaviruses indicates that each of these domains encodes a non-redundant function, which may further facilitate metaviral capsid-RNA interactions or encode other functions to potentially defend metaviral RNAs from host defenses. Thus, by capturing snapshots in time based on when they were domesticated, metaviral-derived host genes provide important archeological insights into metaviral protein domains and retrotransposition strategies. The mysterious N-terminal non-capsid domain of the ARC protein may represent just such an archaeological clue about an ancient domain with no extant homologs that was acquired either from Metaviridae or host genomes ∼350 million years ago.

## Abbreviations

CA: capsid
ERV: endogenous retrovirus
INT: integrase
LTR: long terminal repeat
MA: matrix
My: millions of years
NC: nucleocapsid
ORF: open reading frame
PR: protease
RBD: RNA-binding domain
RT: Reverse transcriptase

## ACKNOWLEDGEMENTS

We thank Caroline Langley, Ching-Ho Chang, Jeremy Hollis, and Peter Dietzen, for comments on the manuscript and for valuable discussions. Research in the Wiedenheft lab is supported by the NIH (R35GM134867), the M.J. Murdock Charitable Trust, a young investigator award from Amgen, the Montana State University Agricultural Experimental Station (USDA NIFA), and a sponsored research agreement from VIRIS Detection Systems. Research in the Malik lab is supported by a subaward from NIH grant U54 AI170792 (PI: Nevan Krogan) and by an Investigator award from the Howard Hughes Medical Institute. Molecular graphics and analyses performed with UCSF ChimeraX, developed by the Resource for Biocomputing, Visualization, and Informatics at the University of California, San Francisco, with support from National Institutes of Health R01-GM129325 and the Office of Cyber Infrastructure and Computational Biology, National Institute of Allergy and Infectious Diseases. We thank Coltran Hophan-Nichols and the rest of Montana State University’s RCI (Research Cyberinfrastructure) team for computational support. Funders had no role in the conceptualization, designing, data collection, analysis, decision to publish, or preparation of the manuscript. We are grateful to numerous genome centers for making assemblies publicly available.

## METHODS

### Building capsid HMMs

To generate capsid-specific Hidden Markov Models (HMMs), we first collected diverse capsid-like sequences for each *Ortervirales* order found in vertebrates (*Retroviridae, Metaviridae, Pseudoviridae,* and *Belpaoviridae)* (Vargiu et al. 2016; Gifford et al. 2018; Krupovic et al. 2018). We began with five previously generated PFAM HMMs (now in Interpro) from the clans Viral_Gag CL0148 (n=5 HMMs) and Gag-polyprotein CL0523 (n=10) (El-Gebali et al. 2019). However, submitting the seed alignment for each HMM to HHPred (Zimmermann et al. 2018) revealed that 13 out of these 15 HMMs in these PFAM clans do not precisely match the capsid domain. Therefore, we built new HMMs that precisely identified the capsid domain in each major clade of retrovirus (n=4 clades) and LTR retrotransposons (n=4 clades). To build custom HMMs for each group of reverse transcribing virus capsid, we queried NCBI’s non-redundant protein database (Pruitt et al. 2007) with PFAM seed alignments (in cases where the HMM covered the full-length capsid domain) or single sequences using a single iteration of PSI-BLAST (Altschul et al. 1997). Sequences identified by PSI-BLAST were aligned in MAFFT (Rozewicki et al. 2019) and the alignment submitted to HHpred to identify the capsid domain using structural homology prediction. Alignments were trimmed to the full-length capsid domain, and pressed into profile HMMs using hmmbuild from the HMMER3 package (Eddy 2011) to generate eight retrovirus or retrotransposon-specific HMMs (**Figure S1A, Table S1**). Each clade of LTR retroelement has a single capsid HMM, except for *Metaviridae*. When our initial analysis did not identify a strong hit for ARC, we built a second metaviral HMM based off of previously generated ARC structures (Hallin et al. 2018).

To verify the specificity of capsid HMMs, a single sequence from each class of LTR retroelement was queried with each of the eight HMMs constructed. Each HMM identified a single domain from only one of these control sequences with a high bit score and a highly significant E-value, while returning low-scoring hits (in the case of ARC/Metaviridae HMM) or no hit at all for capsid sequences from other LTR retroelement clades. In sequences associated with previously determined structures (Ball et al. 2016; Qu et al. 2018; Acton et al. 2019; Nielsen et al. 2019), HMM matches overlap exactly with the bi-lobed alpha-helical capsid domains. In sequences with no known molecular structure, AlphaFold (Jumper et al. 2021) was used to predict tertiary structure. In these unstudied sequences, HMMs identified domains for which AlphaFold predicted bi-lobed alpha-helical architecture (**Figure S1B-C**).

### Identifying capsid genes in vertebrate genomes

To generate a comprehensive catalog of capsid-like sequences in the human genome, we generated a six-open reading frame translation of the recently complete T2T genome (GCF_009914755) (Nurk et al. 2021; Rhie et al. 2022) using the EMBOSS (Rice et al. 2000) tool getorf with default parameters (-find = 2 (between START and STOP codons), minimum size 75bp), identifying 127,070,530 ORFs. ORFs were queried for capsid homology using the previously described HMMs, using hmmscan with default parameters and without gathering or length thresholds. We repeated our search on 25 additional vertebrate genomes downloaded from UCSF or NCBI to identify as many metaviral capsid domains as possible for our subsequent phylogeny of metaviral capsid domains. Additionally, we downloaded Repbase 25.03 (Bao et al. 2015) and generated a six-frame translation of all sequences in Repbase to identify previously characterized consensus sequences for LTR retroelements. This approach could be less sensitive to capsid domains interrupted by large insertions or introns; however, a similar search of predicted spliced transcripts found in RefSeq yielded no additional hits. Furthermore, previously annotated capsid-containing domesticated genes have no introns within the capsid region (Campillos et al. 2006; Kokošar and Kordiš 2013).

### Matching human capsid-like ORF sequences to previously annotated genes

To understand whether human HMM matches were part of existing gene annotations, or represented previously uncharacterized sequences, we used the sequence of each match as a blastn query against NCBI’s RefSeq database, considering only exact matches. We obtained the full-length open reading frames of the matching annotated genes for use in our subsequent analyses. This approach allowed us to identify many previously unidentified orthologs across placental mammals (**Figure 2**).

### Analysis of domain architecture and genetic context

To identify other metaviral fragments (protein domains or LTRs) that could remain in or adjacent to each human domesticated metaviral capsid genes, we extracted flanking nucleic acid sequences 16 kb upstream of the start codon and 16 kb downstream of the stop codon from the T2T genome assembly. These ∼33 kb genomic sequences were queried for long terminal repeats (LTRs) using LTRHarvest (Ellinghaus et al. 2008). We generated new six-frame translations of the ∼33 kb extracted sequences using getorf (-find=1 between STOP codons and -find = 2 between START and STOP codons, with a minimum size of 75bp). We queried translated sequences with the entire PFAM 34.0 database to identify any recognizable protein domains nearby as well as our capsid HMMs to confirm previous capsid identification. These HMM search results allowed us to identify the conserved non-canonical start site of CCDC8 (Figure S9).

### Phylogenetic analysis of capsid-like sequences

To understand the evolutionary relationships between capsid-like sequences in the human genome and between all metaviral capsid sequences found in 26 vertebrate genomes, we built two maximum likelihood phylogenetic trees (**Figure 1**, **Figure 3**). Full-length capsid sequences were defined as ORFs longer than 150 amino acids, containing HMM alignments longer than 125 amino acids (**Figure S1**). These were aligned using MAFFT (Rozewicki et al. 2019) (using the more accurate L-ins-I method). Alignments were trimmed and filtered using trimAl (gt = 0.5)(Capella-Gutiérrez et al. 2009) and manually inspected for sequences that remained with internal insertions and deletions. A phylogenetic tree was then constructed in Fasttree2 (Price et al. 2010) using the JTT evolutionary model with gamma-shape rate variation based on ProtTest model selection (Abascal et al. 2005). Phylogenetic trees were visualized with the ggtree package in R (Yu et al. 2017). Tree topologies were verified by building phylogenetic trees in IQTREE (Minh et al. 2020) using both UFBOOT and traditional bootstrap parameters. We found that metaviral sequences consistently separated from endogenous retroviral sequences in the human phylogeny. Mammalian metaviral orthologs consistently separated out into four unique clades with unique groups of non-mammalian metaviral sequences as the closest-branching outgroup.

### Identification of mammalian orthologs

To identify potential orthologs of the human domesticated metaviral-derived capsid-like genes, we performed thorough searches of 17 additional representative mammalian genomes (14 placental mammals, two marsupials, and one monotreme) using several complementary methods. Our process was iterative, combining multiple rounds of database searches followed by sequence analysis (alignment building and orthology checking via phylogenies).

We initiated the first search using the full-length human ORF nucleotide sequences and their translations as initial queries for tblastn searches (protein query versus nucleotide database) of predicted transcripts in the NCBI non-redundant nucleotide database, specifying the species of interest for each search. In subsequent search step iterations, we used our initial alignments and phylogenies (see below) to identify species/gene combinations where more attention was needed, either because a gene seemed to be missing, or because the gene prediction appeared to be truncated. To search for these ‘missing’ genes, we employed additional strategies that are better designed to detect degenerate pseudogenes. We added blastn searches (nucleotide query versus nucleotide database), we searched genome assemblies rather than predicted gene sets In some cases, we added queries from non-human species, choosing closely related species to the one with the missing gene. For some of the apparent pseudogenes, we added sequences from sister taxa (outside the original 18 mammalian species selected) to investigate whether inactivating mutations are real rather than genome assembly errors: seeing inactivating mutations shared across related species gives us greatly increased confidence they are real mutations. We complemented these blast search results by searching six-frame genome translations using our two metaviral HMMs (as we did for human), ensuring that all matches were included in our final sequence collection.

For the analysis phase of these iterations, we generated alignments and phylogenies to help us determine orthologous relationships (depicted in **Figure 2**) as well as to understand which sequences are intact genes versus pseudogenes. Blastn searches against a database of all human transcripts were also a useful tool in this step, often immediately clarifying that a query sequence obviously grouped with a single human gene to the exclusion of all others. For example, using each human metaviral-derived sequence as a blast query against all human genes revealed the very high similarity between genes in the PNMA6E/F, PNMA6A/B and RTL8A/B/C groups, indicating that these human genes fall into only three clades, rather than one for each individual human gene. Using these initial blastn-based group assignments, we generated ‘master’ in-frame nucleotide alignments, using the MACSE frame-aware alignment tool (Ranwez et al. 2018), alignments that we added to and refined during subsequent search/analysis iterations.

Analyzing these in-frame nucleotide alignments using a custom script allowed us to classify each sequence as intact (complete ORF), incomplete due to genome assembly gaps (*i.e.,* we observed one or more N bases and/or the start/end of an assembly contig near the homologous region) or a clear pseudogene (*i.e.,* contains frameshift and/or premature stop codons with respect to the full-length annotated human ORF,or is significantly truncated without evidence of assembly gaps). Our script makes an exception for the *PEG10* gene, because it has been shown to contain a programmed ribosomal frameshift; thus, we permitted frameshifts observed within 10 codons of the expected location.

During our analyses, we noticed that NCBI’s predicted transcripts are sometimes annotated to indicate that potential frameshifting/ premature stop codons that were observed in genome assemblies have been ‘corrected’, with the idea they most likely represent genome sequencing errors. While this approach makes sense for genes that are rarely pseudogenized, we found that for some of these domesticated capsid genes, it may underestimate the number of true pseudogeness. For this reason, we ensure that we always used corresponding sequences extracted directly from genome assemblies, rather than the predicted transcripts in the NR database. For most apparent pseudogenes, we were able to gain additional confidence that inactivating mutations are real by adding sequences from closely related taxa, where we observed that identical inactivating mutations are often shared between taxa.

In order to more formally confirm orthology relationships, we selected full-length intact genes from all species, generated amino acid translations, and realigned them using MAFFT (Rozewicki et al. 2019). Because sequence divergence is very high between the major clades shown in Figure 3A, we aligned sequences only within each of those clades. We manually trimmed each clade’s alignments to contain only well-aligning regions, used ProtTest to select the best-fitting evolutionary model, and used PHYML (Guindon et al. 2010) to generate phylogenies, with 100 bootstrap replicates (JTT or LG substitution model, --pinv e --alpha e -f e). Visual inspection of these trees allowed us to confidently assign the overarching orthology relationships. To confirm orthology assignments for pseudogenes and recent duplicates, we also generated nucleotide-based phylogenies of the small in-frame alignments for individual groups, again using PHYML (Parameters: GTR substitution model, flags --pinv e --alpha e -f e). We also checked that species relationships roughly followed the expected species tree (**Figure 2**). Finally, we obtained gene age estimates in My (millions of years) using the TimeTree database (Kumar et al. 2022).

### Analysis of evolutionary selective constraints

We performed two complementary analyses of the evolutionary selective pressures acting on these genes. First, we used sequences from diverse placental mammals to estimate the ‘overall’ selective pressure on each gene. From our master in-frame nucleotide alignments for each gene, we selected only the intact genes, from only the 18 placental mammal species shown in Figure 2. We removed any in-frame gaps from the resulting alignments, and manually trimmed each to contain only well-aligning regions. We generated a phylogeny from each trimmed alignment (using PHYML with the GTR substitution model, estimating the proportion of invariable sites, shape of the gamma distribution and nucleotide frequencies. We then used the alignment and tree as input to the codeml algorithm from the PAML package (Yang 2007) to estimate overall dN/dS, using “model 0”, which makes the simplifying assumption that all lineages and all amino acid sites evolve under uniform selective pressure (additional parameters: codon model=2, initial dN/dS = 0.4, cleandata=0). We also ran codeml under model 0 but fixing dN/dS=1 to examine the null hypothesis that sequences evolve neutrally. We determined the level of statistical support for rejecting the null (neutral) hypothesis in favor of the alternative model where dN/dS has the value estimated by model 0 by comparing twice the difference in log-likelihoods of these two model 0 runs to a chi-squared distribution with 1 degree of freedom (**Table 1**, Model 0 (Placentals) column)).

Second, while the overall dN/dS estimates indicate that purifying selection is the prevailing evolutionary pressure operating on these genes, we wanted to explore the hypothesis that positive selection might be acting at a subset of amino acid positions in some of these genes. For this position-specific analysis we used primate genomes, which provide a powerful clade in which to look for selection (the placental mammal alignments contain much greater diversity, such that synonymous substitutions begin to show saturation). We therefore used blastn searches of NCBI’s non-redundant database to collect primate orthologs for each gene, again using reciprocal blastn searches against a local database of all human genes to ensure one-to-one orthology and selecting only intact genes for our analysis. We generated in-frame alignments using MACSE, adjusted and trimmed alignments manually, and then estimated phylogenies using PHYML as before. Again, we used PAML’s codeml algorithm (codon model=2, initial dN/dS = 0.4, cleandata=0), but this time we compared the log likelihoods of evolutionary models that allow for a subset of residues under positive selection (model 8, model 2) with paired models that only allow purifying and neutral selection (models 8a/7, model 1). For genes where maximum likelihood tests indicate positive selection, we report in Table 1 the proportion of sites estimated to under positive selection, the dN/dS of this class of sites, and the identity of sites with a >90% posterior probability of being members of the positively selected class (codeml’s “BEB” Bayes Empirical Bayes method). For these genes, we also checked the robustness of evidence favoring positive selection by re-running analyses with alternative codeml parameters (codon model=3, starting dN/dS=3). We also used GARD (Kosakovsky Pond et al. 2006) to rule out the possibility that genetic recombination may have caused false positive signals of positive selection (Anisimova et al. 2003). Even after analysis with GARD, we found robust evidence for positive selection in multiple genes at multiple sites.

### Structural modeling

To gain structural insight into the regions of human metaviral sequences not annotated by either our capsid HMMs or Pfam, we downloaded all human metaviral sequences from AlphaFold’s Protein Structure Database (Varadi et al. 2022) and submitted unannotated portions of the protein to DALI and Foldseek to identify structural homologues (Holm 2020; Kempen et al. 2022). This strategy identified numerous RNA binding domains with N-terminus was submitted to DALI. For example, the N-terminus of PNMA1 identified 939 significant hits (Z-score > 2) in the PDB, the majority of which were RNA-binding domains. The top ten hits all contained Z-scores above 6.6 for RNA-binding proteins (See supplementary materials for DALI output). AlphaFold was also used to predict the structures of the closest consensus metaviral sequences from Repbase, and structural homology between metaviral sequences was determined using structural superposition (mmaker command in the ChimeraX Daily build) (Pettersen et al. 2021). This strategy confirmed the near perfect retention of capsid domains and revealed that the majority of metaviral-derived genes have regions of predicted disorder with no clear globular domains.

### Expression analysis

Tissue-specific expression data was downloaded from the Genotype-Tissue Expression (GTEx) Project, supported by the Office of the Director of the National Institutes of Health, and by NCI, NHGRI, NHLBI, NIDA, NIMH, and NINDS (Lonsdale et al. 2013). The median gene level TPM dataset used for the expression analysis were obtained from the GTEx Portal on 10/13/21. A publicly available RNA-seq dataset for placental expression data was downloaded from the Human Protein Atlas for placenta-specific expression on October 12, 2021 (https://www.proteinatlas.org/about/download)(Uhlén et al. 2015). A third RNA-seq data set was downloaded from a placenta-specific expression study (Gong et al. 2021) (https://www.obgyn.cam.ac.uk/placentome/), and fragments per kilobase per million (FPKM) values were converted to transcript per million (TPM) values. A heatmap of expression was made in R using the pheatmap package.

### International Mouse Phenotyping Consortium

HGNC gene names were submitted as queries to the IMPC database. Results for genes in which knockout mice had been made were summarized in table format (**Table S5**).

