## Supplemental figures for "The diverse evolutionary histories of domesticated metaviral capsid genes in mammals"

### SUPPLEMENTARY FIGURES AND TABLES

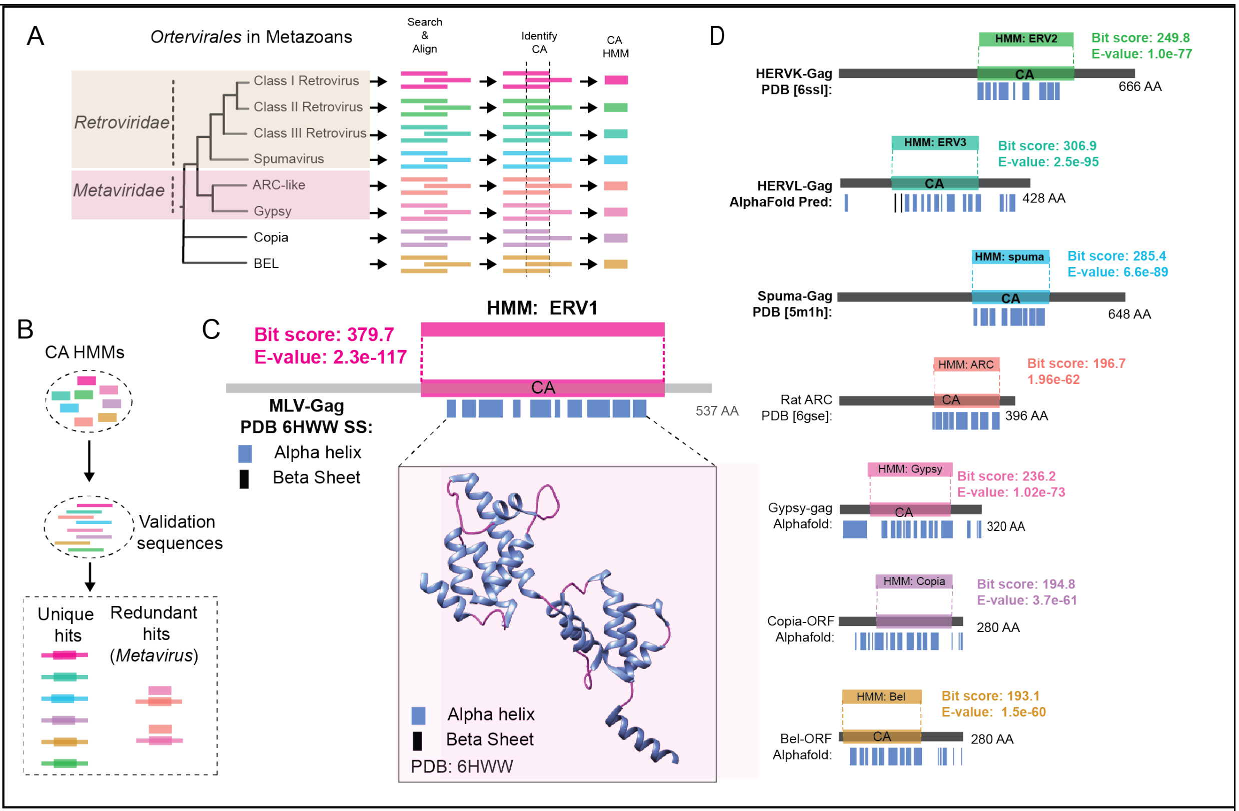


**Figure S1.** Constructing Hidden Markov Models for the capsid domain of LTR retrotransposons. A. Seed sequences from four groups from *Retroviridae*, two groups of *Metaviridae* (Gypsy), and one group each of *Pseudoviridae* (Copia) and *Belpaoviridae* (BEL) were used to generate clade-specific full-length capsid HMMs (schematic phylogeny derived from (Gifford et al. 2018; Krupovic et al. 2018)). B. Capsid HMMs were validated by querying a pool of sequences containing a single sequence from the clades in (A). C-D. Capsid HMMs cover the full-length capsid domain and identify positive control sequences with high bit-scores and low e-values.

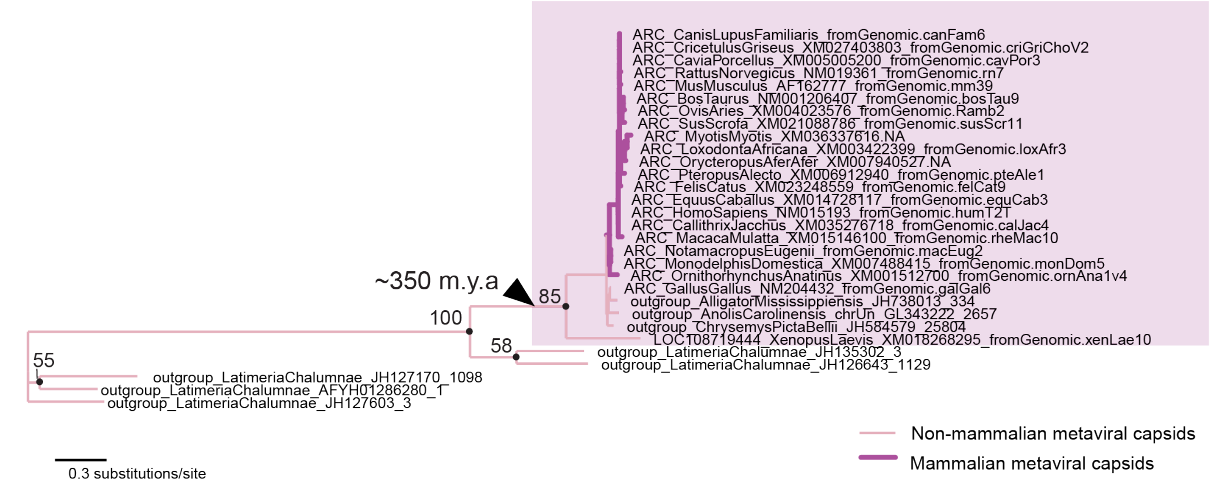


**Figure S2. Phylogeny of ARC proteins**. Arc-like sequences and proxy ancestral metaviruses from coelacanth (*Latimeria chalumnae)* were translated and aligned in MAFFT. Phylogenetic tree was built from an amino acid alignment in PHYML with 100 bootstrap replicates.

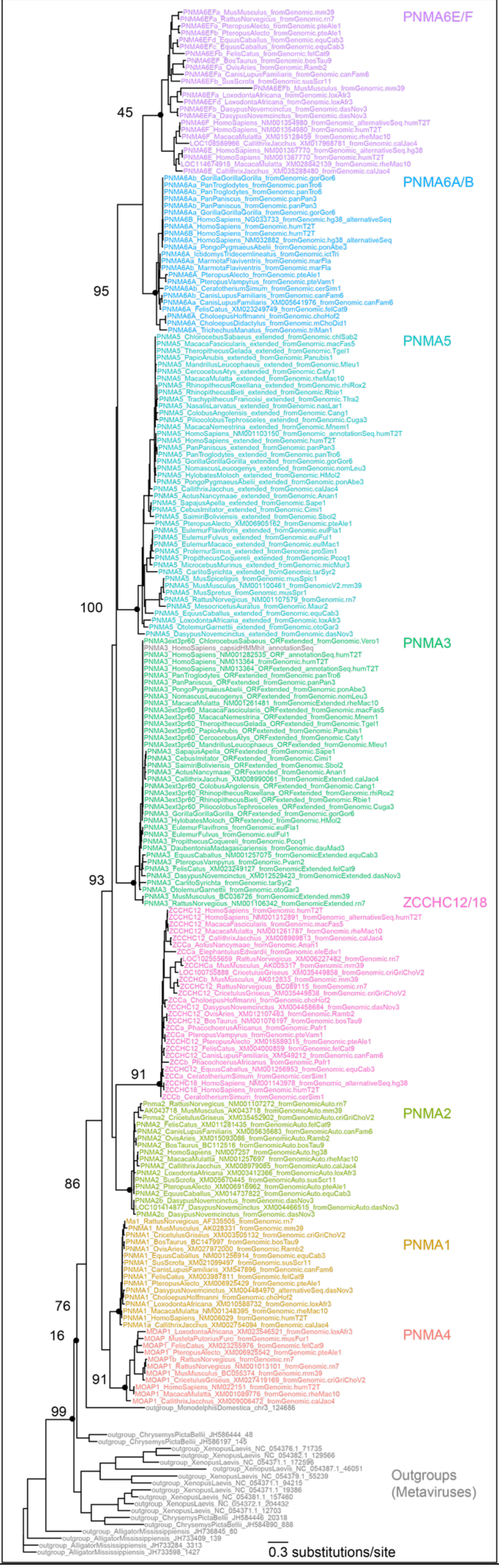
PNMA6E/F
PNMA6A/B
PNMA5
PNMA3
ZCCHC12/18
PNMA2
PNMA1
PNMA4
Outgroups
(Metaviruses)
45
95
100
93
91
86
76
16
91
99
0.3 substitutions/site

**Figure S3. Phylogeny of PNMA proteins.** Full-length intact *PNMA* genes (including the single *PNMA* gene from marsupials) were translated and aligned in MAFFT as were outgroup metaviral sequences from the African clawed frog (*Xenopus laevis*), the painted turtle (*Chrysemys picta belli)* and the American alligator (*Alligator mississippiensis*). Alignments were manually trimmed and then phylogenies generated in PHYML with 100 bootstrap replicates. Numeric values on the left represent bootstrap support of the corresponding nodes on the right (black dots) The PNMA6E/F clade was not well-resolved in this analysis, so it was analyzed separately (Figure S5).

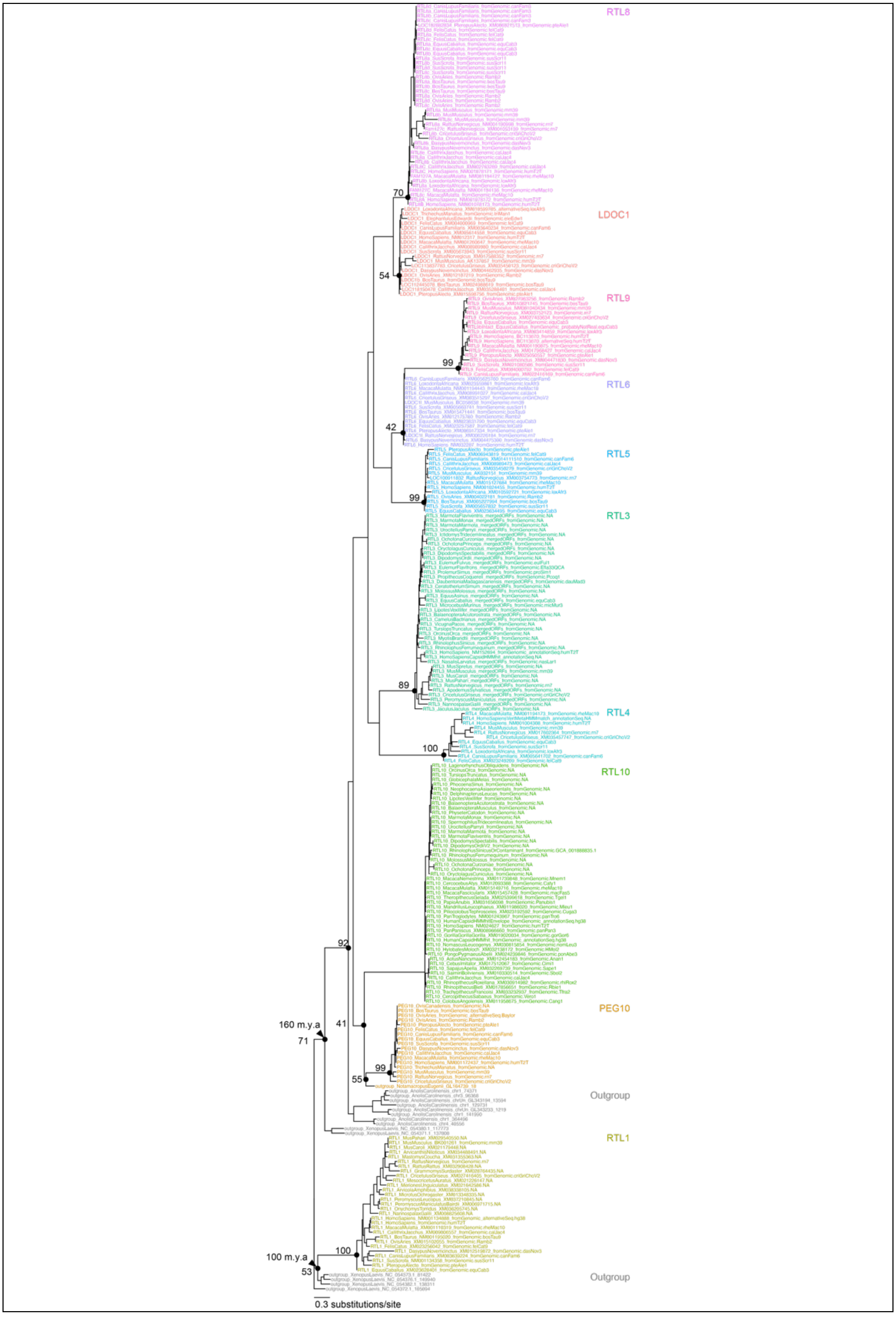

RTL8
70
LDOC1
54
RTL9
99
RTL6
42
RTL5
99
RTL3
89
RTL4
100
RTL10
92
PEG10
160 m.y.a
71
41
99
55
Outgroup
RTL1
100
100 m.y.a
53
Outgroup
0.3 substitutions/site

**Figure S4. Phylogenetic tree for SIRH/RTL and RTL1 proteins**. Full-length intact genes were translated and aligned in MAFFT as were outgroup metaviral sequences from the anole lizard (*Anolis carolensis)* and African clawed frog (*Xenopus laevis*). Alignments were manually trimmed and then phylogenies generated in PHYML with 100 bootstrap replicates. Numeric values on the left represent bootstrap support of the corresponding node on the right (black dots). In this and in other phylogenetic analyses, RTL1 consistently groups distinctly from the other SIRH/RTL proteins, suggesting independent domestication.

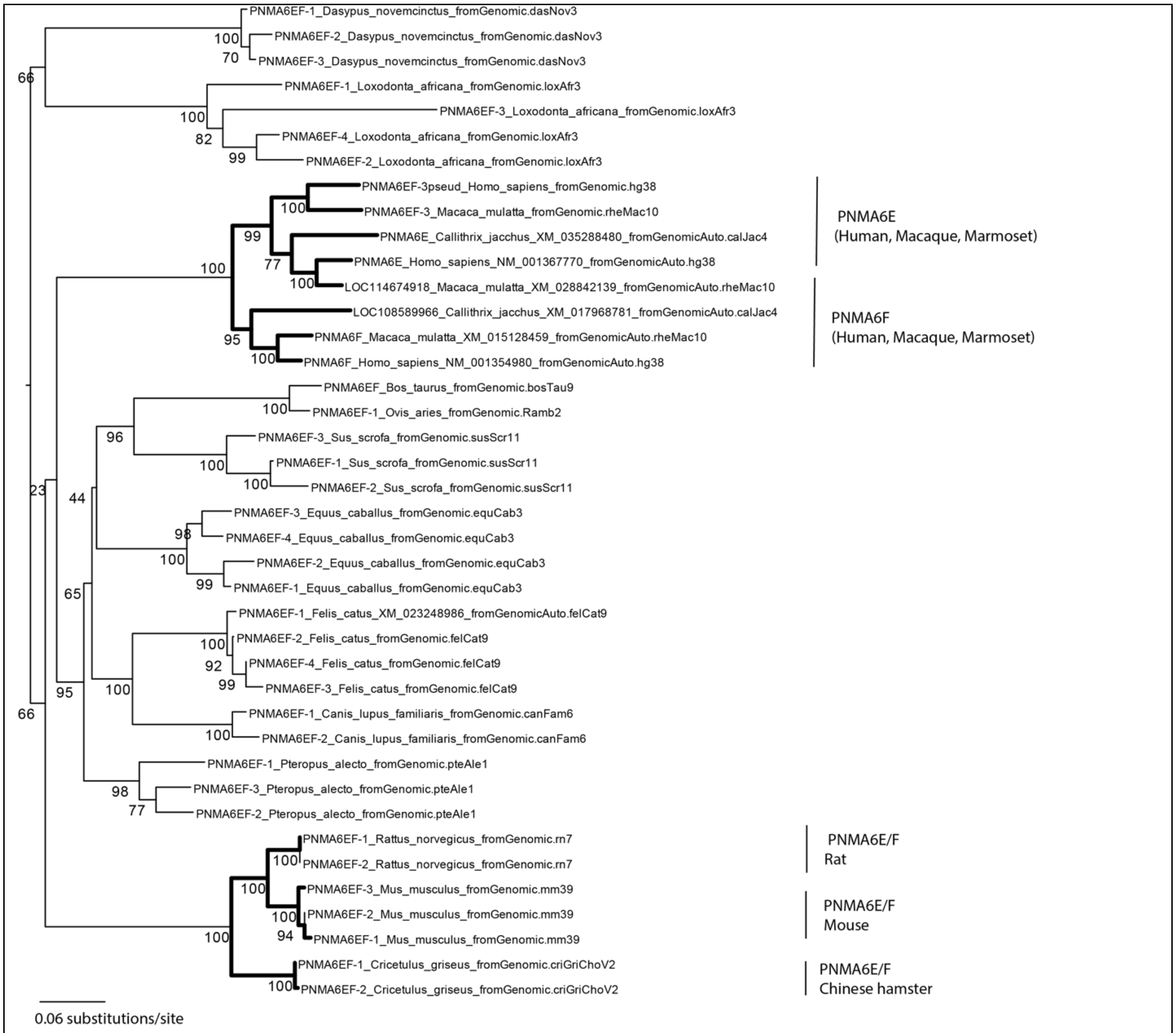


**Figure S5. PNMA6E/F has duplicated multiple times in distinct lineages.** Full-length intact genes were translated and aligned in MAFFT. Alignments were manually trimmed and then phylogenies generated in PHYML with 100 bootstrap replicates. Numeric values on the left represent bootstrap support of the corresponding node on the right (black dots). In several instances PNMA6E/F proteins group with paralogs from the same species, rather than orthologs from other species.

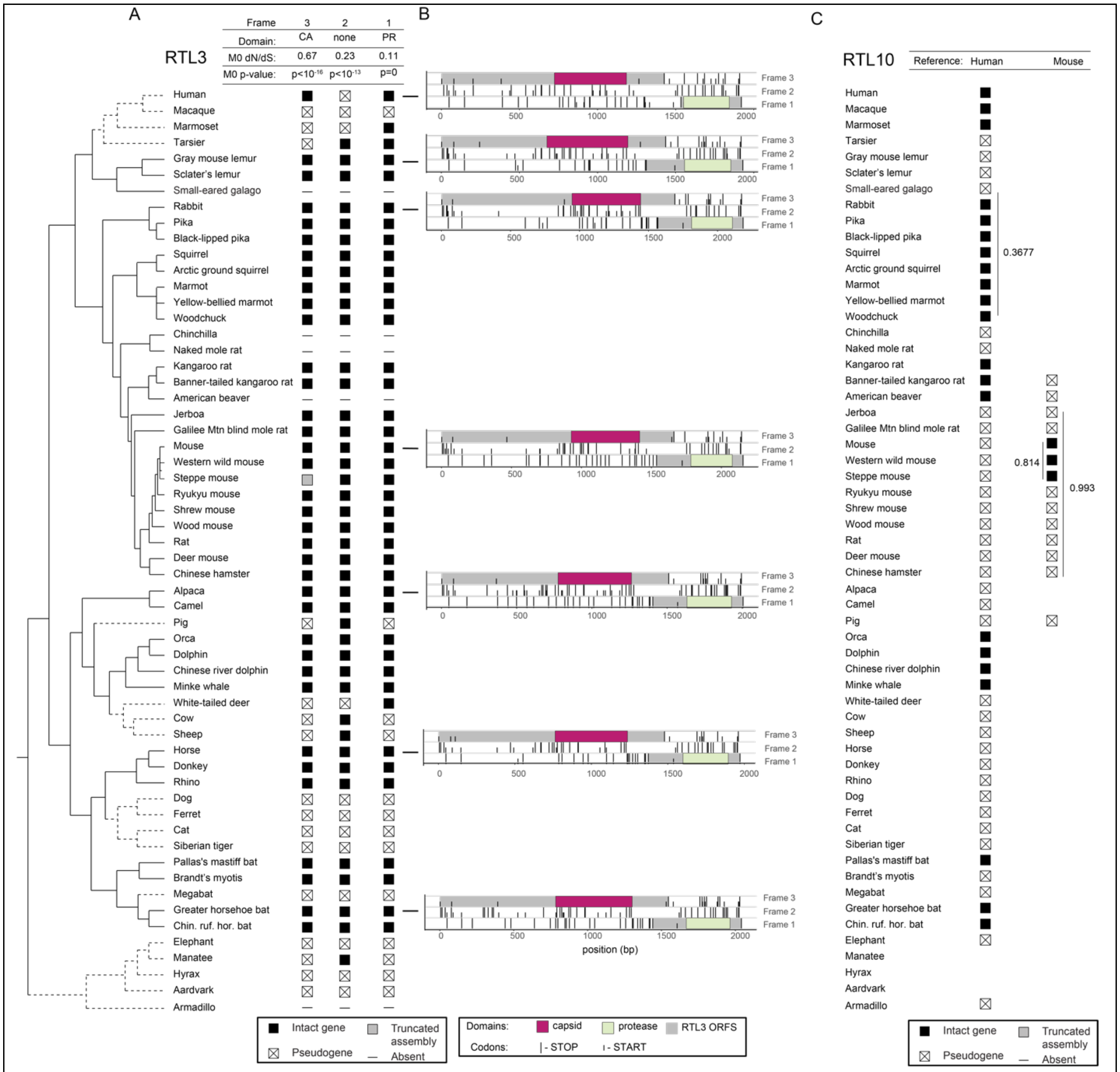


**Figure S6. RTL3 and RTL10 exhibit domain-specific and lineage-specific patterns of conservation**. A. RTL3 is retained in genomes of most placental mammals, but has undergone loss or pseudogenization in several lineages (dotted lines in phylogeny). Filled squares represent intact genes and squares containing a cross represent sequences with obvious inactivating mutations (frameshifts and/or premature stops). Gray boxes represent sequences that are truncated due to genome assembly gaps, and '-' symbols represent cases where we could find no matching sequence. The RTL3 protease domain is more conserved than the capsid domain. B. Three-frame translations of the RTL3 region reveal stop codons that preclude programmed frame-shifts in the "intervening" region between the capsid and protease domains in some lineages; nevertheless, these domains might be independently translated. Long vertical bars indicate stop codons, short vertical bars indicate start codons. Domains other than the capsid and protease domain are colored grey for ease of visualization. C. RTL10 is ancient but has undergone widespread pseudogenization in mammals. However, RTL10 remains well conserved and under purifying selection in several lineages, including rodents (dN/dS calculated from codeml Model 0 are indicated).

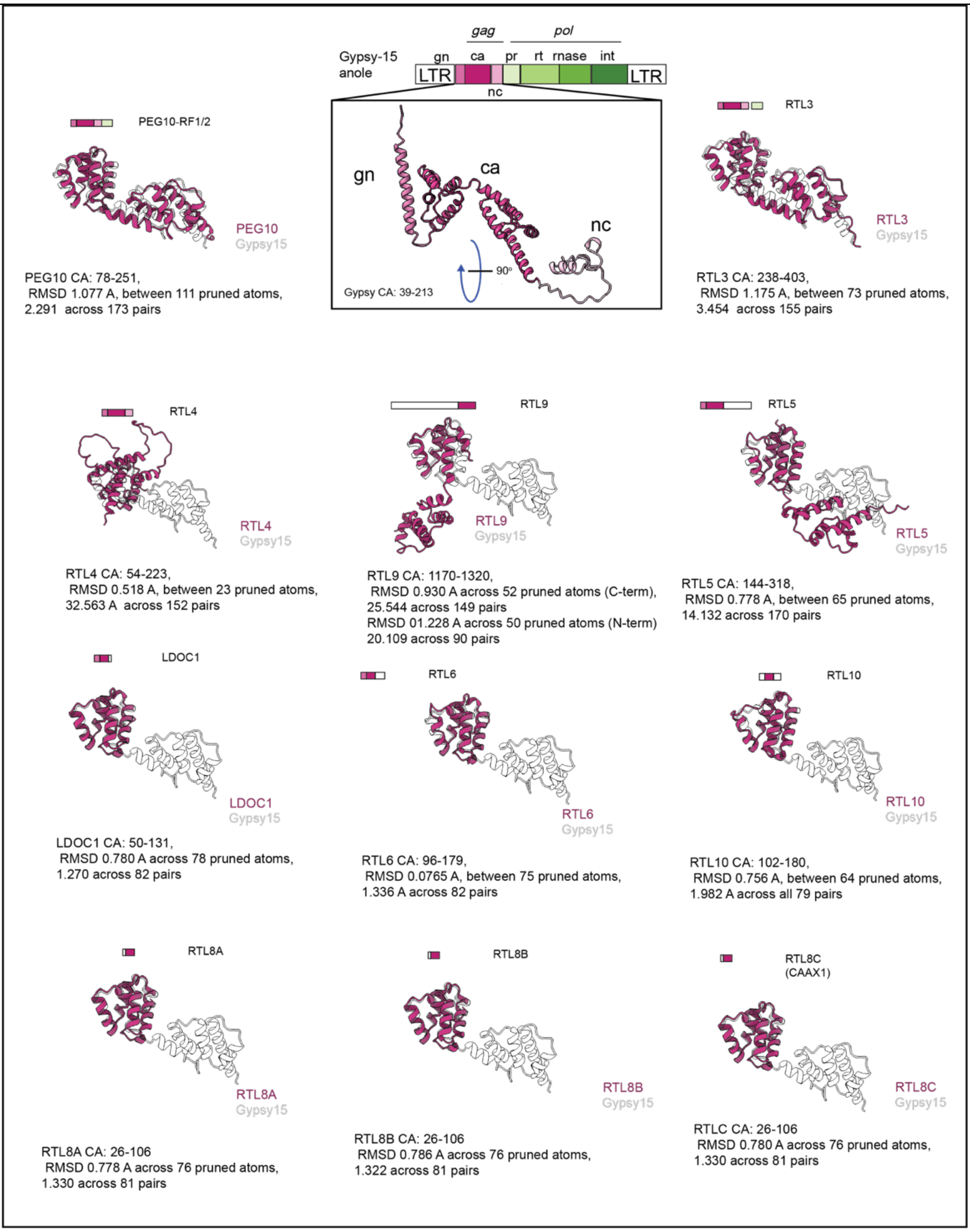


**Figure S7. Conservation and divergence of the capsid domain in the SIRH/RTL family.** We compared the Alphafold predicted structure of each human protein in the SIRH/RTL family to the Alphafold predicted structure of a proxy ancestral metavirus, Gypsy-15 from the anole lizard. Capsid domain coordinates identified by HMMER were aligned to the Gypsy-15 capsid domain in ChimeraX with the *mmaker* command. The N-terminal lobe of the capsid domain is structurally identical across all proteins encoded by the *SIRH/RTL* family of genes. In contrast to the N-terminal lobe of the capsid domain, the C-terminal lobe is only present in PEG10, RTL3, RTL4, RTL9, and RTL5.

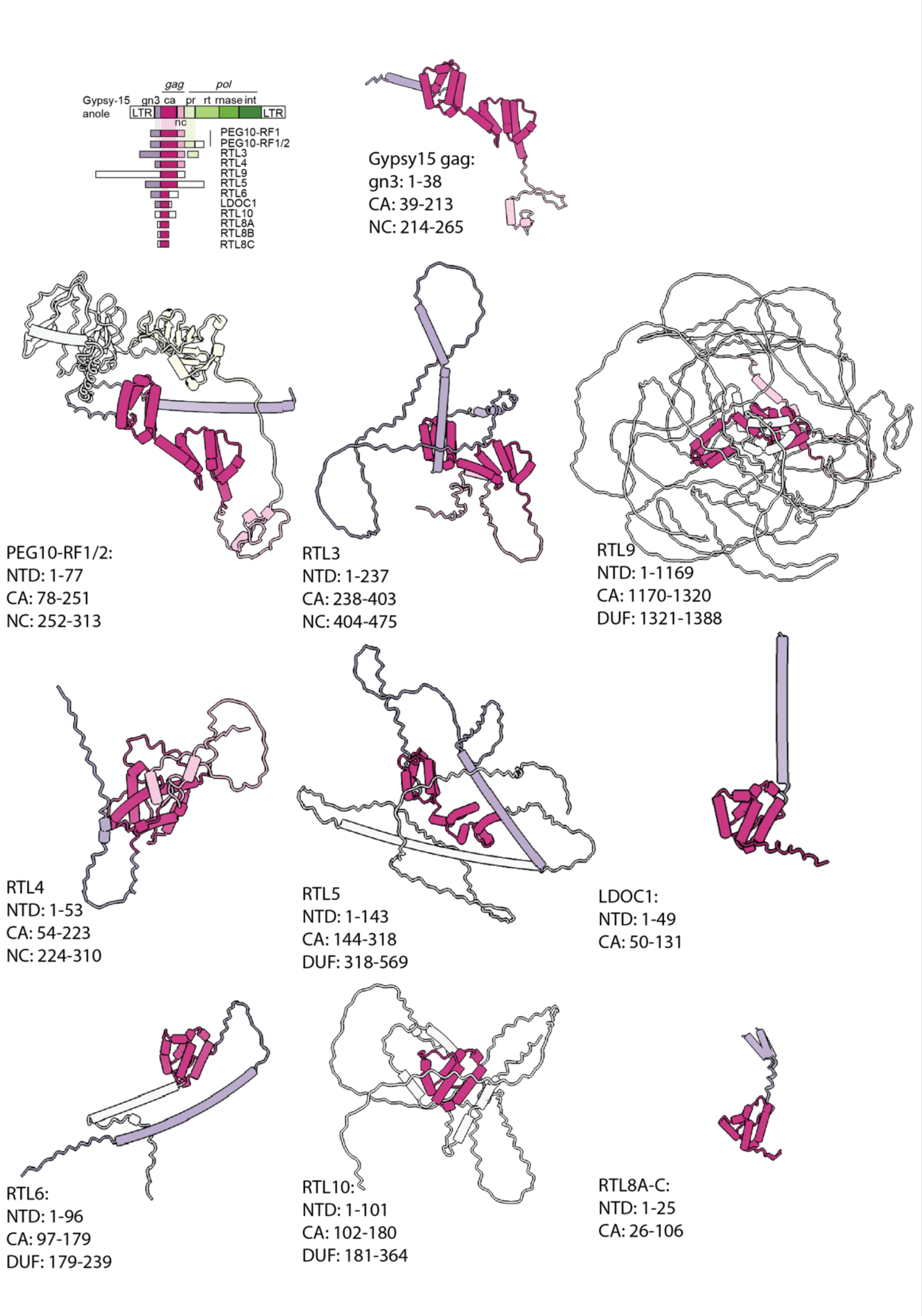


**Figure S8. Lack of conservation in the N-terminal domain SIRH/RTL family**. We compared the length and architecture of Alphafold-predicted structures for the N-terminal domain (NTD) upstream of the capsid (CA) domain (magenta) of the SIRH/RTL family of genes (amino-acid coordinates for each domain listed below each structure). The N-terminal domain is made up of single alpha helices and disordered regions of varying lengths (light purple) and lacks conservation across the SIRH/RTL gene family.

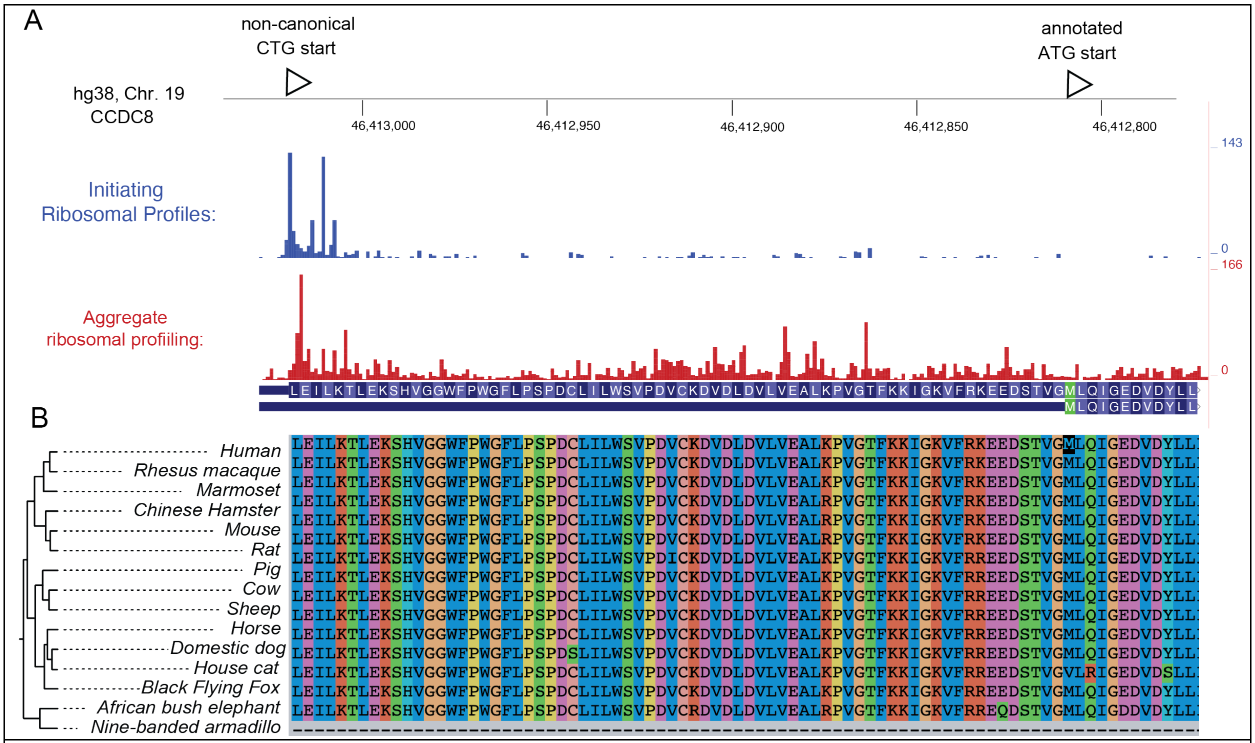


**Figure S9. CCDC8 contains a conserved non-canonical translation start site that encodes a full-length PNMA-family RNA-binding domain**. A. Aggregated ribosomal profiling studies from the UCSC genome browser show an accumulation of initiating ribosomes at the non-canonical CTG start site. B. The non-canonical start site is highly conserved across placental mammals. As a result, CCDC8 encodes an intact and conserved RNA-binding domain (RBD) similar to other RBDs in the PNMA family (**Figure 4**).

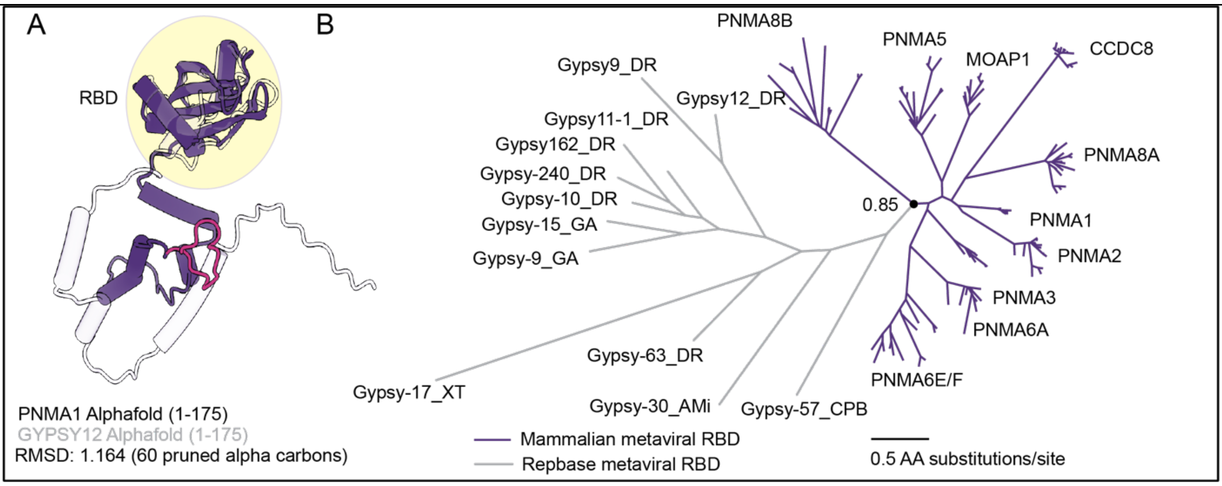


**Figure S10. The evolutionary origins of the RNA-binding domain of the PNMA family**. The RNA-binding domain (RBD) of the PNMA family originated from metaviruses in a single domestication event. The RBD is still found in actively circulating *Metaviridae***.** A. Structural overlay of the first 100 residues of Alphafold predictions of a metaviral and PNMA1 RBDs shows a high degree of structural conservation (RMSD calculated from *mmaker* in ChimeraX). B. Mammalian and metaviral RBDs form distinct clades. Amino acid sequences identified by an RBD-specific HMM were aligned and a maximum-likelihood phylogenetic tree built in FastTree (maximum-likelihood support for the monophyletic origin of the PNMA RBDs is indicated with a black dot).

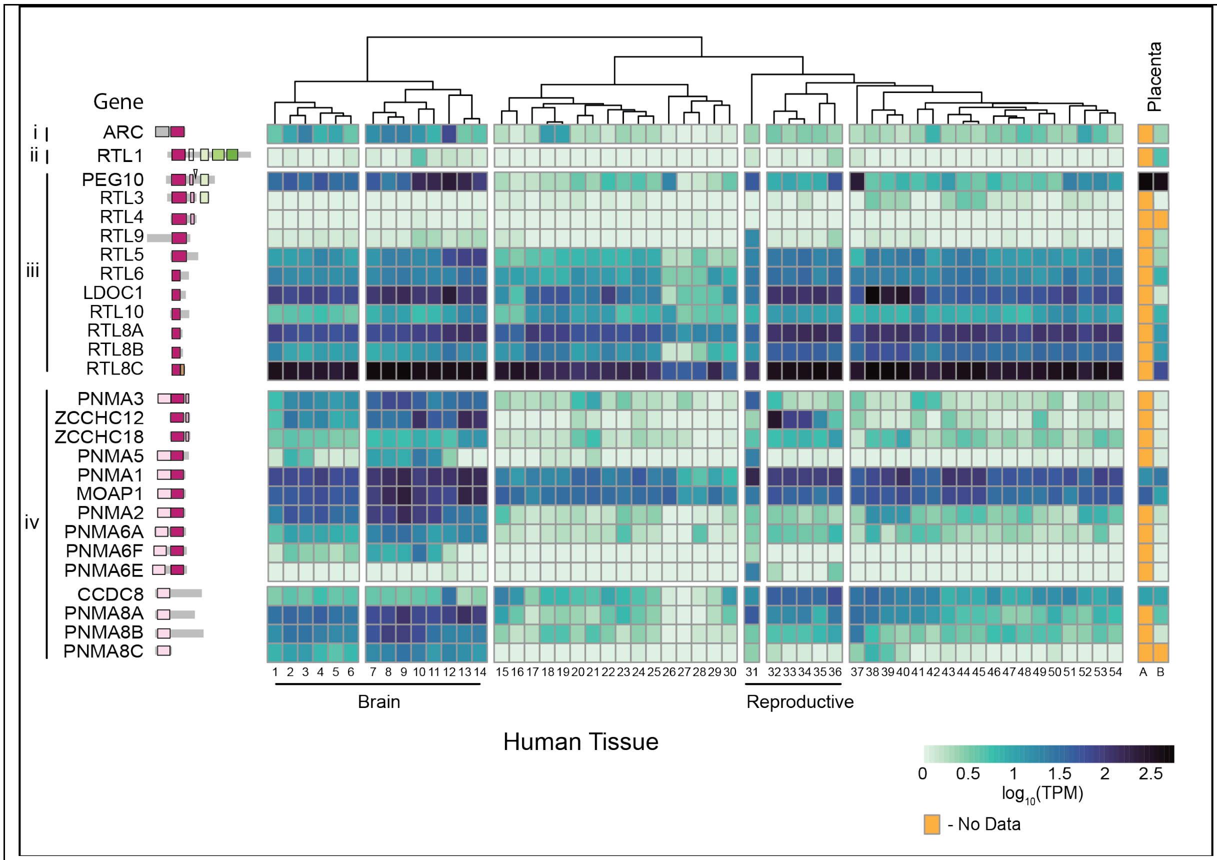


**Figure S11. Human gene expression of domesticated metaviral capsid genes**. Bulk RNA-sequencing data for 54 human tissues was downloaded from the GTEX portal, and median transcript per million (TPM) value plotted for each gene in each tissue. Data from the placenta were downloaded from the Human Protein Atlas and from Gong *et al.* and converted to median TPM values. Expression data reveals that most domesticated metaviral genes are widely expressed in human tissues, while some are only expressed in specific tissues (e.g. *RTL1* in the brain and placenta).
