## Supplementary material for "The diverse evolutionary histories of domesticated metaviral capsid genes in mammals": Table S1

**Table S1.** Model statistics for custom capsid HMMs. Bit-scores and E-values are for the control sequence from each clade used to test the HMMs.

| **LTR Retroelement** | **MSA (n)** | **MSA (length)** | **HMM length** | **Bit-score** | **E-value** |
| --- | --- | --- | --- | --- | --- |
| ***Retroviridae* - Class I ERV** | 149 | 294 | 249 | 379.7 | 2.30E-117 |
| ***Retroviridae* - Class II ERV** | 173 | 270 | 213 | 249.8 | 1.00E-77 |
| ***Retroviridae* - Class III ERV** | 17 | 203 | 197 | 306.9 | 2.50E-95 |
| ***Retroviridae* - Spumavirus** | 14 | 175 | 175 | 285.4 | 6.60E-89 |
| ***Metaviridae* - Gypsy** | 96 | 220 | 192 | 236.2 | 1.02E-73 |
| ***Metaviridae* – ARC** | 55 | 197 | 149 | 196.7 | 1.96E-62 |
| ***Pseudoviridae* - Copia** | 212 | 207 | 174 | 194.8 | 3.70E-61 |
| ***Belpaoviridae* - BEL** | 221 | 204 | 180 | 193.1 | 1.50E-60 |
