## Supplementary material for "The diverse evolutionary histories of domesticated metaviral capsid genes in mammals": Table S3

**Table S3.** BLAST statistics for consensus Repbase metaviral sequences used as proxies for ancestral metaviruses. Translated N-terminal domains from the start codon to the start of the capsid domain were aligned and scored using pBLAST.

|  | Gypsy-4 | Gypsy-15 | Gypsy 2-1 |  |
| --- | --- | --- | --- | --- |
| Gypsy-4 |  |  |  |  |
| 1-68 |  |  |  |  |
| Gypsy-15 | -33 |  |  | Pairwise BLAST score |
| 1-28 | 8% |  |  | Percent ID |
|  | 21% |  |  | Percent similarity |
| Gypsy 2-1 | -123 | -155 |  | Pairwise BLAST score |
| 1-196 | 8% | 6% |  | Percent ID |
|  | 17% | 7% |  | Percent similarity |
| Gypsy-30 | -47 | -70 | -100 | Pairwise BLAST score |
| 1-100 | 15% | 8% | 12% | Percent ID |
|  | 28% | 12% | 23% | Percent similarity |
