## Supplementary material for "The diverse evolutionary histories of domesticated metaviral capsid genes in mammals": TAble S5

**Table S5.** Phenotypes associated with knockouts of domesticated metaviral genes in mice. P-values for knockouts from the International Mouse Phenotyping Consortium of domesticated metaviral genes are reported for each observed phenotype.

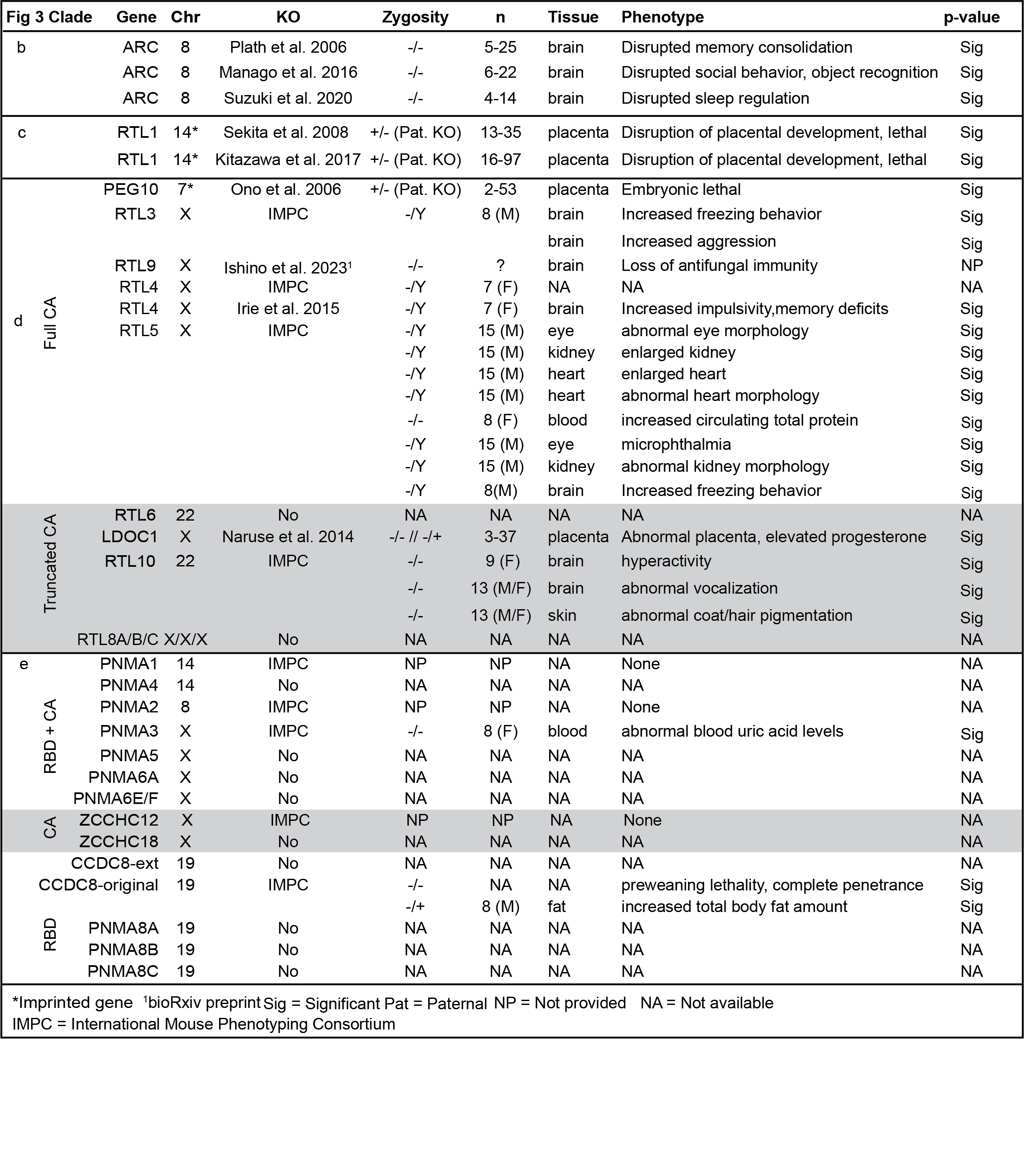
